# A scalable and robust variance components method reveals insights into the architecture of gene-environment interactions underlying complex traits

**DOI:** 10.1101/2023.12.12.571316

**Authors:** Ali Pazokitoroudi, Andrew Dahl, Noah Zaitlen, Saharon Rosset, Sriram Sankararaman

**Affiliations:** Department of Computer Science, UCLA, Los Angeles, CA, USA; Department of Human Genetics, David Geffen School of Medicine, UCLA, Los Angeles, CA, USA; Department of Computational Medicine, David Geffen School of Medicine, UCLA, Los Angeles, CA, USA; Department of Neurology, UCLA, Los Angeles, CA, USA; Section of Genetic Medicine, Department of Medicine, University of Chicago, Chicago, Illinois, USA; Department of Statistics, Tel-Aviv University, Tel-Aviv, Israel

## Abstract

Understanding the contribution of gene-environment interactions (GxE) to complex trait variation can provide insights into mechanisms underlying disease risk, explain sources of heritability, and improve the accuracy of genetic risk prediction. While biobanks that collect genetic and deep phenotypic data over large numbers of individuals offer the promise of obtaining novel insights into GxE, our understanding of the architecture of GxE in complex traits remains limited. We introduce a method that can estimate the proportion of trait variance explained by GxE (GxE heritability) and additive genetic effects (additive heritability) across the genome and within specific genomic annotations. We show that our method is accurate in simulations and computationally efficient for biobank-scale datasets.

We applied our method to *≈* 500, 000 common array SNPs (MAF *≥* 1%), fifty quantitative traits, and four environmental variables (smoking, sex, age, and statin usage) measured across *≈* 300, 000 unrelated white British individuals in the UK Biobank. We found 69 trait-environmental variable pairs with significant genome-wide GxE heritability (*p <* 0.05*/*200 correcting for the number of trait-E pairs tested) with an average ratio of GxE to additive heritability *≈* 6.8% that include BMI with smoking (ratio of GxE to additive heritability = 6.3 *±* 1.1%), WHR (waist-to-hip ratio adjusted for BMI) with sex (ratio = 19.6 *±* 2%), LDL cholesterol with age (ratio = 9.8 *±* 3.9%), and HbA1c with statin usage (ratio = 11 *±* 2%). Analyzing nearly 8 million common and low-frequency imputed SNPs (MAF *≥* 0.1%), we document an increase in genome-wide GxE heritability of about 28% on average over array SNPs. We partitioned GxE heritability across minor allele frequency (MAF) and local linkage disequilibrium values (LD score) of each SNP to observe that analogous to the relationship that has been observed for additive allelic effects, the magnitude of GxE allelic effects tends to increase with decreasing MAF and LD. Testing whether GxE heritability is enriched around genes that are highly expressed in specific tissues, we find significant tissue-specific enrichments that include brain-specific enrichment for BMI and Basal Metabolic Rate in the context of smoking, adipose-specific enrichment for WHR in the context of sex, and cardiovascular tissue-specific enrichment for total cholesterol in the context of age. Our analyses provide detailed insights into the architecture of GxE underlying complex traits.

## Introduction

Variation in a complex trait is modulated by an interplay between genetic and environmental factors. Characterizing the effects of gene-environment interactions (GxE) on complex trait variation has the potential to shed light on biological mechanisms underlying the trait [1, 2, 3], inform public health measures [4], identify sources of missing heritability [5], and to improve the accuracy and portability of trait prediction [6, 7]. The growth of biobanks that collect genetic and deep phenotypic data (that span disease outcomes, clinical labs, lifestyle factors, and environmental exposures) across large numbers of individuals offers the possibility to gain novel insights into GxE [8, 3]. Nevertheless, characterizing GxE has proved challenging due, in part, to the small effect sizes of individual genetic variants [9, 10].

A potentially powerful methodological approach aims to quantify GxE effects aggregated across a set of variants without needing to pinpoint individual variants. In this approach, the proportion of trait variation explained by GxE (GxE heritability or 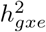 ) is estimated by fitting a class of variance components models where the model parameters, *i*.*e*., the variance components, are informative of 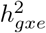. Methods for estimating 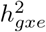 using this approach include GCTA-GxE [11], multitrait GREML (MV-GREML) [5], random regression GREML (RR-GREML) [5, 12], whole-genome reaction norm model (RNM) and its multitrait version (MRNM) [13]. All of these methods (except RNM) are able to account for differences in the noise or residual variance across environments (noise heterogeneity), which is important to mitigate biases in GxE heritability estimates [13, 14]. However, these methods work with discrete-valued environmental variables with RNM and MRNM further restricted to fit bivariate and univariate environments, respectively. A more recent general framework, GxEMM [14] can be applied to both discrete and continuous environmental variables while modeling noise heterogeneity. However, none of these methods are practical for biobank-scale datasets with sample sizes in the hundreds of thousands and genetic variants in the millions. Two recent methods, GPLEMMA [15] and MEMMA [16], attempt to scale GxE heritability estimation to large-scale datasets but do not model noise heterogeneity. As a result, current methods for estimating GxE heritability either do not scale to the biobank setting or are susceptible to biased estimates. Additional insights into the architecture of GxE can be gleaned if we can move beyond genome-wide estimates of GxE heritability and estimate GxE heritability across specific genomic annotations such as common vs. low-frequency SNPs or functional genomic annotations.

We propose a scalable and robust method, GENIE (**G**ene-**EN**vironment **I**nteraction **E**stimator) that can estimate the proportion of trait variance explained by GxE and additive genetic effects (additive heritability). Using extensive simulations and real data analysis, we show that GENIE accurately estimates 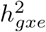 and provides calibrated tests of 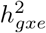 due to its ability to account for noise that is heterogeneous across environments. Importantly, GENIE is scalable: able to estimate GxE on datasets with hundreds of thousands of individuals, millions of SNPs, and tens of environmental variables in several hours. The ability of GENIE to be applied to large-scale datasets is important for power: we show that GENIE has adequate power to detect 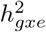 as low as 2% across a sample of *≈* 300 K unrelated individuals. Finally, GENIE is versatile: able to handle multiple environmental variables (discrete or continuous) and to estimate not only genome-wide 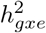 but also partition 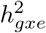 across genomic annotations (both overlapping and non-overlapping). To demonstrate its utility, we first applied GENIE to estimate the genome-wide 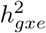 on common SNPs (*M* = 459, 792 SNPs with MAF *>* 1%) and four environmental variables (smoking, sex, age, and statin usage) for fifty quantitative phenotypes measured across 291, 273 unrelated white British individuals in the UK Biobank (UKBB). Second, we leveraged the scalability of GENIE to partition 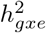 across common and low-frequency imputed SNPs (*M* = 7, 774, 235 with *MAF >* 0.1%) in UKBB. We partitioned 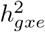 into genomic annotations based on the minor allele frequency (MAF) and local linkage disequilibrium (LD score) of each SNP to investigate the variation in GxE effects with population genetic features and to estimate genome-wide 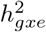 that includes the contribution of both common and low-frequency SNPs. Finally, we applied GENIE to assess whether 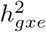 shows tissue-specific enrichment by analyzing each of 53 tissue-specific gene sets identified from the GTEx dataset [17].

## Results

### Methods overview

GENIE aims to jointly estimate the proportion of phenotypic variance that can be explained by genetic (G), gene-environment interactions (GxE), and noise-environment interactions or noise heterogeneity (NxE) by fitting a variance component model that relates phenotypes ***y*** measured across *N* individuals to their genotypes ***X*** (across *N* individuals *× M* SNPs), environmental covariates ***E*** (where each column corresponds to one of *L* environmental covariates), and fixed-effect covariates ***C*** as follows:

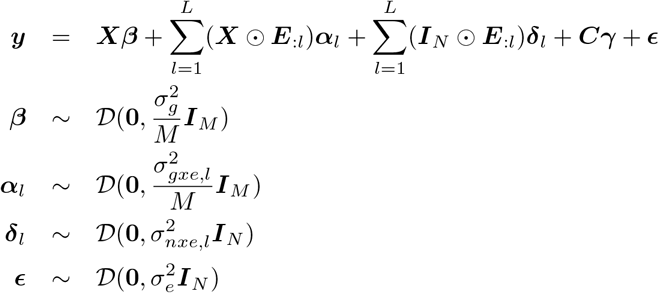

Here *D*(***µ*, Σ**) is an arbitrary distribution with mean ***µ*** and covariance **Σ. *E***_:*l*_ denotes column *l* of ***E***, and ⊙denotes row-wise Kronecker product. ***β*** denotes the *M* -vector of SNP effect sizes, ***α***_*l*_ denotes the *M* -vector of genetic effect sizes in the context of environment *l* (GxE effects), ***δ***_*l*_ denotes the *N* -vector of NxE effect sizes for environment *l*, and ***E*** denotes the *N* -vector of noise. In this model, 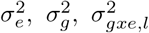, and 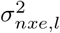 are the variance components associated with noise, additive genetic effects, gene-by-environment effects for a given environment, and noise-by-environment effects for a given environment respectively. These variance components can then be transformed into the additive heritability or the proportion of variance explained by additive effects (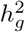 associated with 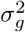) and the GxE heritability or the proportion of variance explained by interactions of genetics with a given environment (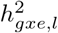 associated with 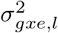 ). We include the environmental covariates as fixed effects following prior work that has shown this approach to ameliorate biases due to gene-environment (G-E) correlation [13, 14] (although G-E correlation is unlikely to bias estimates of polygenic GxE that are being estimated here as shown in [14]).

The model underlying GENIE is flexible. The ability to model noise-environment interactions allows for different residual or noise variance across values of the environmental covariates. This increased flexibility can control false positive inferences of GxE heritability. Further, GENIE allows for jointly estimating the proportion of phenotypic variance due to genetic interactions with single or multiple environmental variables. Finally, GENIE can also partition GxE heritability across both non-overlapping and overlapping genomic annotations or components using the following model (Methods).

The key challenge in applying this model to biobank-scale datasets is that estimation of variance components is computationally prohibitive. To overcome this challenge, GENIE uses a randomized method-of-moments estimator of the variance components wherein the algorithm works with a sketch of the genotype matrix obtained by multiplying the genotype matrix with a chosen number of random vectors *B*. When the number of random vectors *B* is small, the resulting algorithm has both scalable runtime and memory usage (Methods).

### Calibration and Power

We assess the false positive rate of tests of GxE heritability based on GENIE in simulations under different genetic architectures with no GxE heritability. For each architecture, we simulated 100 phenotype replicates across *N* = 291, 273 unrelated white British individuals in the UKBB and *M* = 459, 792 SNPs with MAF *>* 1% genotyped on the UK Biobank genotyping array. We chose statin usage in the UKBB as the environmental variable. We varied the percentage of causal SNPs while fixing the additive heritability at 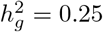. We ran GENIE with *B* = 10 random vectors (see Section on Effect of the choice of the number of random vectors).

Across all simulations, the false positive rate of rejecting the null hypothesis of no GxE heritability is controlled at levels 0.05 and 0.05*/*200 (we consider this threshold which controls for the number of trait-E pairs that we test in UKBB): the average *P* (rejection at *p < t*) is 4% and 0% for *t* = 0.05 and *t* = 0.05*/*200 respectively (Figure 1a)).

**Figure 1:**
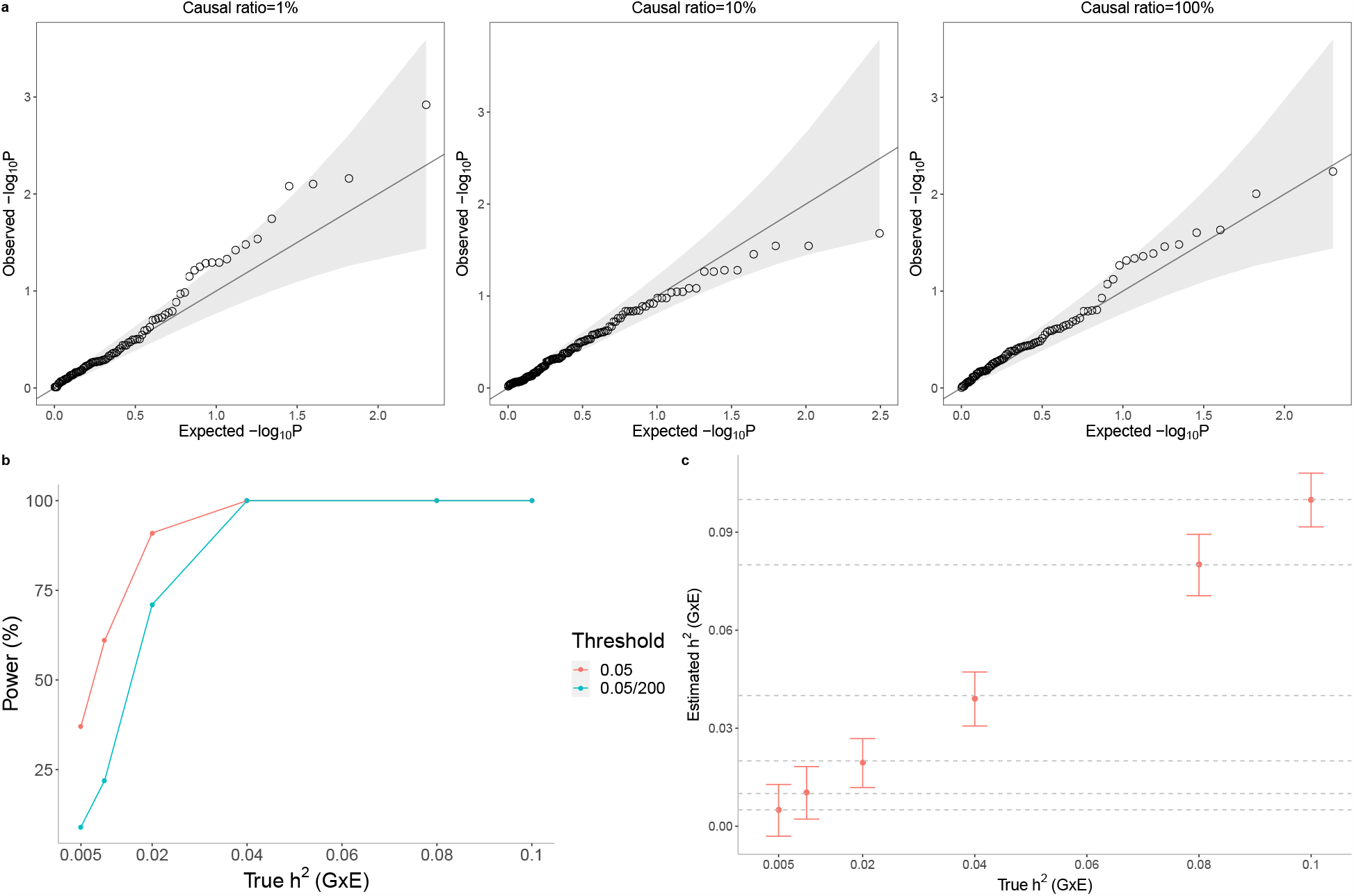
Calibration and power of GENIE in simulations (*N* = 291, 273 unrelated individuals, *M* = 459, 792 SNPs). **a**) Q-Q plot of p-values (of a test of the null hypothesis of zero GxE heritability) when GENIE is applied to phenotypes simulated in the absence of GxE effects. Each panel contains 100 replicate phenotypes simulated with additive heritability 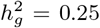 and varying proportions of causal variants. Across all architectures, the mean of *P* (rejection at *p < t*) are 7.5% and 0% for *t* = 0.05 and 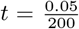 respectively (7.5% is not significantly different from the nominal rate of 5%; the p-value of a test of bias of point estimates of 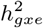 is *p* = 0.75). **b**) The power of GENIE across genetic architectures as a function of GxE heritability. We report power for p-value thresholds of 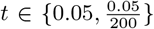. **c**) The accuracy of 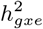 estimates obtained by GENIE. Across all simulations, statin usage in UKBB was used as the environmental variable.

To measure the power of GENIE to detect GxE heritability, we simulated phenotypes with a non-zero GxE heritability. Across genetic architectures, we varied the GxE heritability while fixing the additive heritability at 0.25 and the percentage of causal SNPs at 10% (these are the default parameters of our simulations unless otherwise specified). We simulated 100 replicates for every genetic architecture. Let 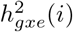 be the estimate of 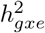 and *SE*_*i*_ be the jackknife estimate of the standard error on the *i*-the replicate for *i ∈ {*1, .., 100*}*. We computed the p-value of a test of the null hypothesis of no 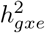 on the *i*-th replicate from the Z-score defined as 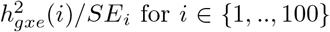. We reported the percentage of replicates with p-value*< t* as the power of GENIE on a given genetic architecture for a p-value threshold of *t*.

GENIE has adequate power to detect GxE effects with 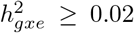 in a sample of 300*K* unrelated individuals at *p <* 0.05 (Figure 1b)). Additionally, across all genetic architectures, GENIE yields unbiased estimates of GxE heritability (Figure 1c)).

Next, we assessed the accuracy of GENIE in a setting where we have multiple environmental variables. We simulated phenotypes from a sub-sampled set of UKBB genotypes choosing a subset of *N* = 10, 000 individuals and 20, 000 SNPs on chromosome 1 of the UK Biobank Axiom array. We considered a setting with *L* = 10 environmental variables with 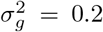, five environmental variables with 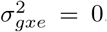, three environmental variables with 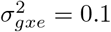and two with 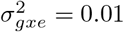. We generated 100 replicates of simulated phenotypes for each set of parameters. We find that GENIE obtains estimates of 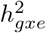 that are accurate across the environmental variables (Supplementary Figure S1, Supplementary Table S1).

### Effect of the choice of the number of random vectors

We explored the choice of the number of random vectors in two ways. First, we quantified the contribution of randomization to the SE of the GxE estimator in GENIE. We simulated 100 phenotypes where 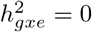. We compared the SE of GxE estimates with *B* = 10 random vectors run 100 times over one of the replicates (the contribution of the randomization to the SE) to the SE of GxE estimates across 100 replicates to determine that, with *B* = 10, randomization contributes to about a third of the total SE. Second, we verified that our GxE estimates are highly correlated for the choice of random vectors *B* = 10 vs *B* = 100 (Pearson’s *ρ* = 0.97; Supplementary Figure S2). These results lead us to conclude that *B* = 10 random vectors provide stable estimates and we use this setting in our remaining analyses.

### Noise heterogeneity

Previous studies have shown that accounting for noise heterogeneity (NxE component) is essential to avoid false positives and inflation in estimates of GxE effects [18, 13, 14]. To demonstrate the importance of modeling NxE, we simulated phenotypes in the presence of NxE effect such that 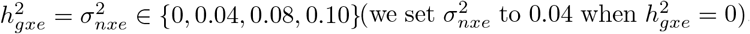. We ran GENIE, in turn, with and without NxE component. Across all simulations, the model that does not account for the NxE component (G+GxE) yields statistically significant upward bias in its GxE estimates (relative bias ranges from 2.5% to 69% across genetic architectures) while the model that fits a noise heterogeneity component (G+GxE+NxE) achieves unbiased estimates of GxE (Figure 2a)).

**Figure 2:**
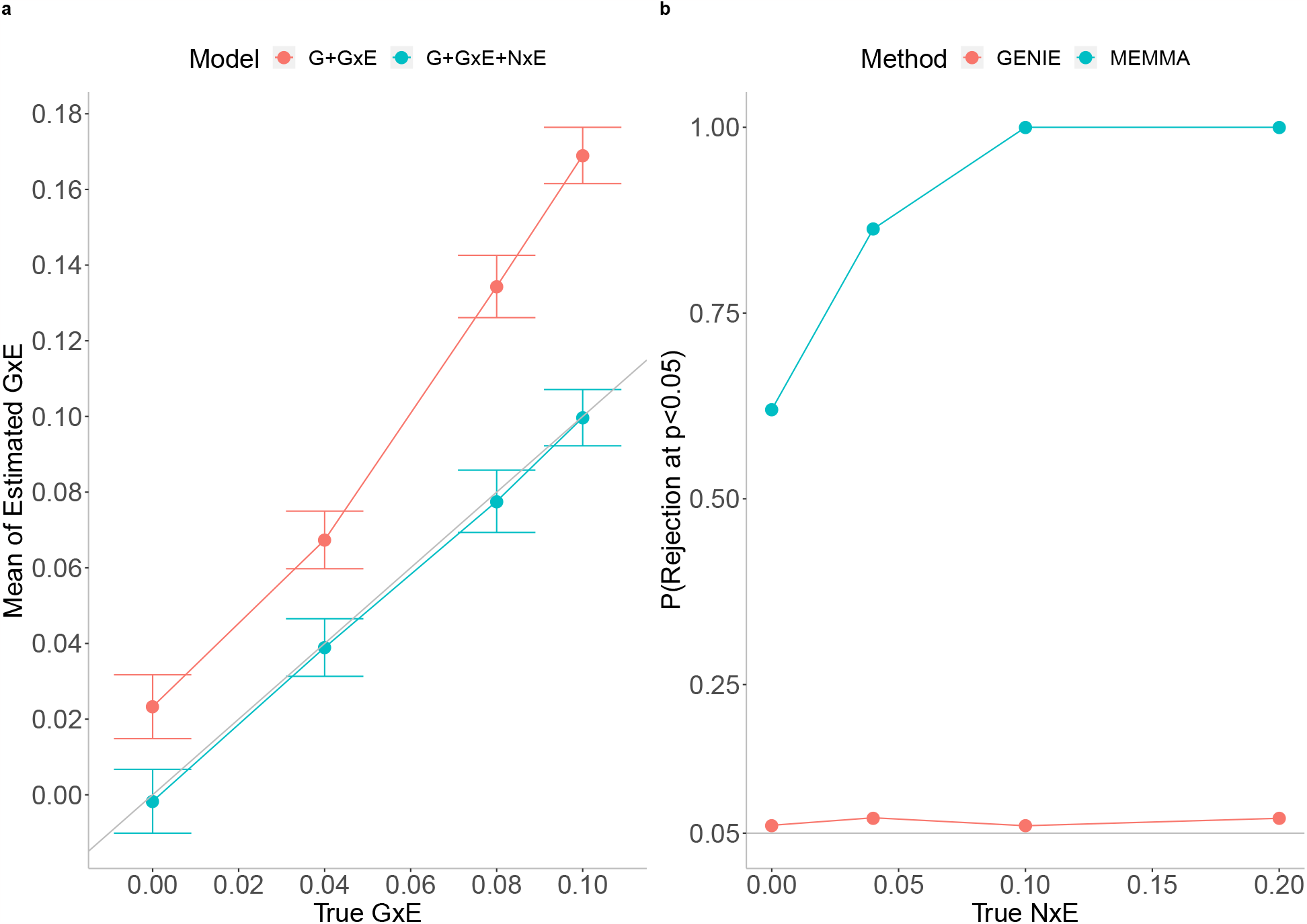
Effect of noise heterogeneity (NxE) on the accuracy of estimates of GxE heritability in simulations. **a**) Comparison of GxE heritability estimates from GENIE under a G+GxE model to those from a G+GxE+NxE model. Model G+GxE refers to a model with additive and gene-by-environment interaction components. Model G+GxE+NxE refers to a model with additive, gene-by-environment interaction, and noise heterogeneity (noise-by-environment interaction) components. We simulated phenotypes with NxE effects and GxE effects across *N* = 291, 273 individuals genotyped at *M* = 459, 792 SNPs. The x-axis and y-axis correspond to the true GxE and the mean of the estimated GxE (from 100 replicates), respectively. Points and error bars represent the mean and *±* SE, respectively. **b**) Comparison of false positive rates of tests for GxE heritability across GENIE and MEMMA. We performed simulations with no GxE heritability but with varying magnitudes of the variance of the NxE effect. We compute the false positive rate as the fraction of rejections (p-value of a test of the null hypothesis of zero GxE heritability *<* 0.05) over 100 replicates of phenotypes simulated from *N* = 40, 000 individuals genotyped at *M* = 459, 792 SNPs.

Further, we compared the calibration of tests of GxE from GENIE with MEMMA [15], a recently proposed scalable method for GxE heritability estimation (we did not include GPLEMMA in this comparison as the model underlying GPLEMMA aims to infer a linear combination of multiple environmental variables that maximizes 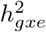 while our experiments all focus on the setting of a single environmental variable). First, we simulated phenotypes with neither GxE effects nor NxE effects from a subset of *N* = 40*k* unrelated white British individuals. In this setting, MEMMA has an inflated false positive rate while GENIE is calibrated (Figure 2b)). The inflated false positive rate for MEMMA in the absence of the NxE effect can be explained by a bias in their estimates of the SE of the variance components (Supplementary Figure S3). We then explored the setting with noise heterogeneity but no GxE. The false positive rate of MEMMA increases as it does not model noise heterogeneity while GENIE has a controlled false positive rate across all simulations (Figure 2b)).

### Computational efficiency

We evaluated the runtime of GENIE, MEMMA, and GCTA(HE) (which implements an exact method-of-moments estimator) with increasing sample size (*N ∈ {*10*K*, 50*K*, 100*K*, 290*K}*) for a fixed number of SNPs (*M* = 459, 792) and a single environmental variable. We ran all methods to fit a single G and GxE variance component. All methods were run on an Intel(R) Xeon(R) Gold 6140 CPU 2.30GHz, with 187GB RAM. GENIE and MEMMA were run with ten random vectors. The runtime of GCTA(HE) includes the time to compute the GRMs. We ran GENIE and GCTA(HE) on a single core while we ran MEMMA on both a single core and four cores. We set a maximum time limit of two days as a constraint for all methods. We could run GCTA(HE) on a dataset of up to 50*K* samples. GENIE is highly scalable and can estimate GxE on about 300*K* individuals and roughly 500*K* SNPs within an hour, approximately 30 times faster than MEMMA run on four cores (Supplementary Figure S4).

### Estimating GxE in the UK Biobank

We applied GENIE to estimate additive heritability 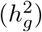 and GxE heritability 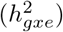 for fifty quantitative phenotypes measured in UKBB across unrelated white British individuals. These fifty phenotypes fall into eight broader phenotypic categories (blood biochemistry, kidney biomarkers, anthropometry, lipid metabolism biomarkers, blood pressure, liver biomarkers, lung, and glucose metabolism biomarkers) that have been analyzed in prior works [19, 20, 21]. Following these studies, we applied a rank-based inverse normal transformation to all phenotypes. We considered, in turn, smoking status, sex, age, and statin usage as environmental variables. We included each environmental variable as a fixed effect in the relevant analyses. First, we explored the importance of modeling noise-environment interactions (NxE) in real data (building on our simulation results). We then analyzed, in turn, common SNPs genotyped on the UK Biobank array (MAF*>* 1%), and then common and low-frequency imputed SNPs (MAF *≥* 0.1%). For select combinations of phenotypes and environmental variables, we also applied GENIE to partition GxE heritability across MAF-LD annotations and to estimate GxE heritability in genes expressed in specific tissues.

#### Robustness of GENIE in the UK Biobank

We first assessed the robustness of GENIE by estimating 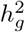 under three different models: G, G+GxE, and G+GxE+NxE where each model is named by the set of variance components fitted jointly. The additive heritability estimates were highly correlated across the models (Pearson’s correlation *ρ ≥* 0.98 for every pair of models), leading us to conclude that GENIE provides robust estimates of additive heritability across different models (Supplementary Figure S5). We observe a significant difference in 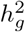 for a handful of trait-E pairs when estimated with G+GxE and G+GxE+NxE that include alcohol frequency intake, overall health, and hair color with both smoking status and sex as environmental variables, and alcohol frequency intake and overall health with age.

Our simulations in the previous section revealed the importance of modeling noise heterogeneity (Figure 2). To investigate the consequences of modeling NxE in real data, we fit, in turn, models without and with NxE (in addition to G and GxE components). The number of trait-E pairs with significant 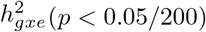 (*p <* 0.05*/*200) decreased from 135 under the G+GxE model to 69 under the G+GxE+NxE model: decreasing from 40 to 21 for smoking (Figure 3b)), 27 to 29 for sex (Supplementary Figure S6b)), 28 to 12 for age (Supplementary Figure S7b)), and 40 to 7 for statin usage (Supplementary Figure S8b)). For traits with significant 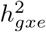, the magnitudes of the estimates varied across the two models: ratio of 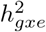 estimates under the G+GxE+NxE to the G+GxE model were 137% on average (range: 43 *−* 350%), 110% (70 *−* 224%), 131% (99 *−* 166%), and 42% (21 *−* 72%) for smoking (Figure 3a)), sex (Supplementary Figure S6a)), age (Supplementary Figure S7a)), and statin (Supplementary Figure S8a)) respectively. The magnitude of noise heterogeneity across trait-E pairs can be substantial: 0.05%, 164%, 10%, and 14% of the additive heritability on average for smoking, sex, age, and statin, respectively (Supplementary Figures S9, S10, S11, and S12).

**Figure 3:**
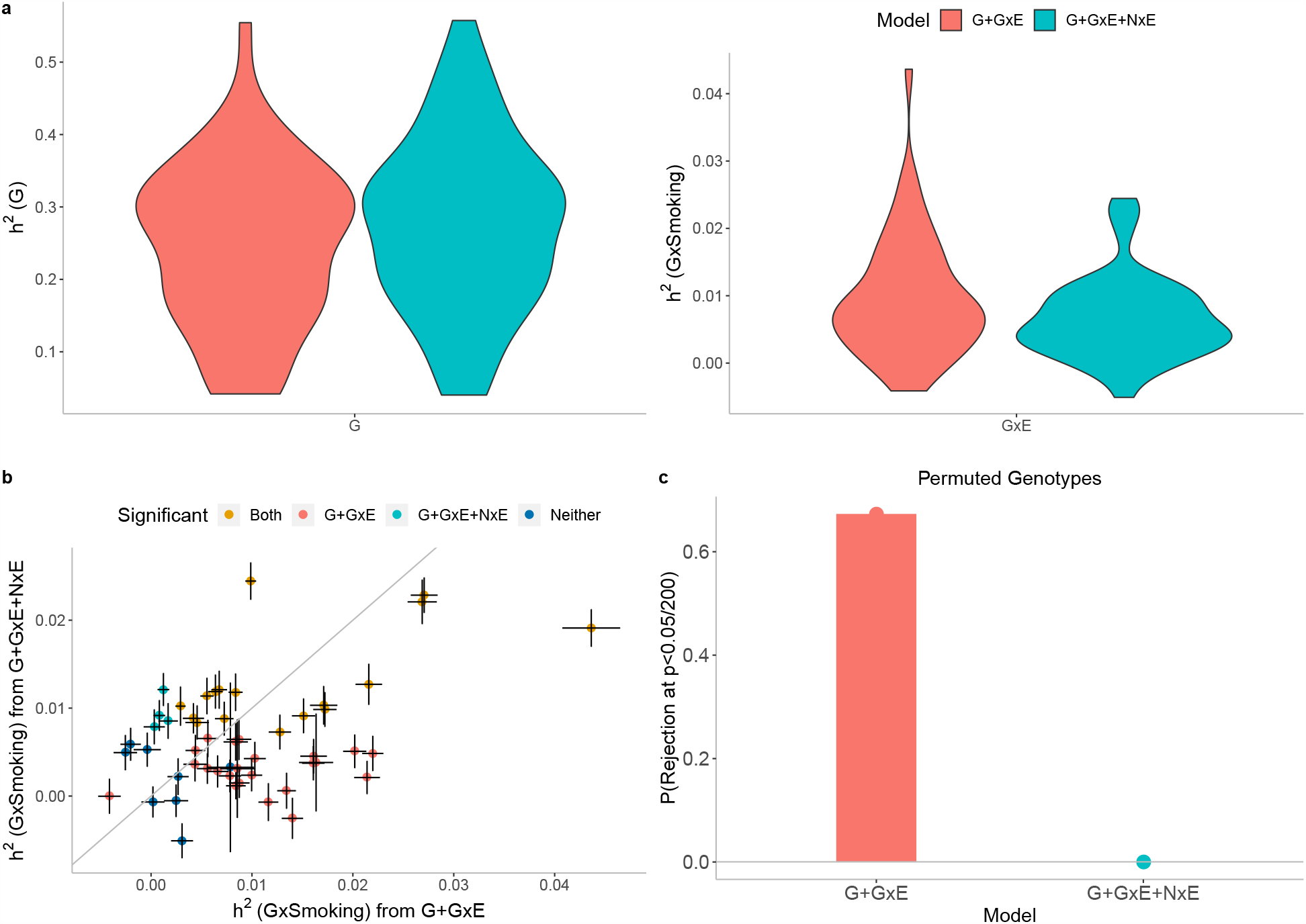
Effect of Noise heterogeneity (NxE) on estimates of heritability associated with GxSmoking across 50 quantitative phenotypes in UKBB. Model G+GxE refers to a model with additive and gene-by-environment interaction components where the environmental variable is smoking status. Model G+GxE+NxE refers to a model with additive, gene-by-environment interaction, and environmental heterogeneity (noise-by-environment interaction ) components. **a**) We run GENIE under G+GxE and G+GxE+NxE models to assess the effect of fitting an NxE component on the additive and GxE heritability estimates. **b**) Comparison of GxE heritability estimates obtained from GENIE under a G+GxE+NxE model (x-axis) to a G+GxE model (y-axis). Black error bars mark *±* standard errors centered on the estimated GxE heritability. Color of the dots indicate whether estimates of GxE heritability are significant under each model. **c**) We performed permutation analyses by randomly shuffling the genotypes while preserving the trait-E relationship and applied GENIE in each setting under G+GxE and G+GxE+NxE models. We report the fraction of rejections (p-value of a test of the null hypothesis of zero GxE heritability 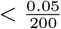 that accounts for the number of phenotypes tested) over 50 UKBB phenotypes.

To further investigate the effect of modeling NxE, we performed permutation analyses by randomly shuffling the genotypes while preserving the trait-E relationship (a setting where there is expected to be no GxE by construction while the relationship between phenotype and E is preserved). We applied GENIE under the G+GxE and G+GxE+NxE models to each trait-E pair. The false positive rate of rejecting the null hypothesis of no GxE across the trait-E pairs is substantially inflated under the G+GxE model while being controlled under the G+GxE+NxE model (Figure 3c), Supplementary Figures S6c), S7c), and S8c) for smoking, sex, age, and statin respectively). These results indicate that modeling NxE is critical to avoid spurious findings of GxE.

#### Gene-by-Smoking Interaction

We applied GENIE to estimate the proportion of phenotypic variance explained by gene-by-smoking in-

teractions 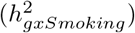 for 50 quantitative phenotypes. We find 21 traits showing statistically significant evidence for 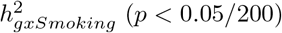 with 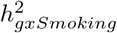 about 6.1% of 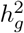 on average (Figure 4). Two of the traits with the largest 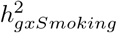 were basal metabolic rate and body mass index (BMI) with estimates of 2.4% and 2.3% respectively (estimates remained significant when we used the binary coding of the smoking status variable obtained by merging the categories of never and previous; Supplementary Figure S13). Our estimates are consistent with a previous study of that analyzed BMI and lifestyle factors in the UKBB to find significant GxE for smoking behavior [5]. The 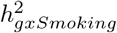 estimates for basal metabolic rate and BMI are about 11% and 7% of their respective 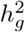 estimates.

**Figure 4:**
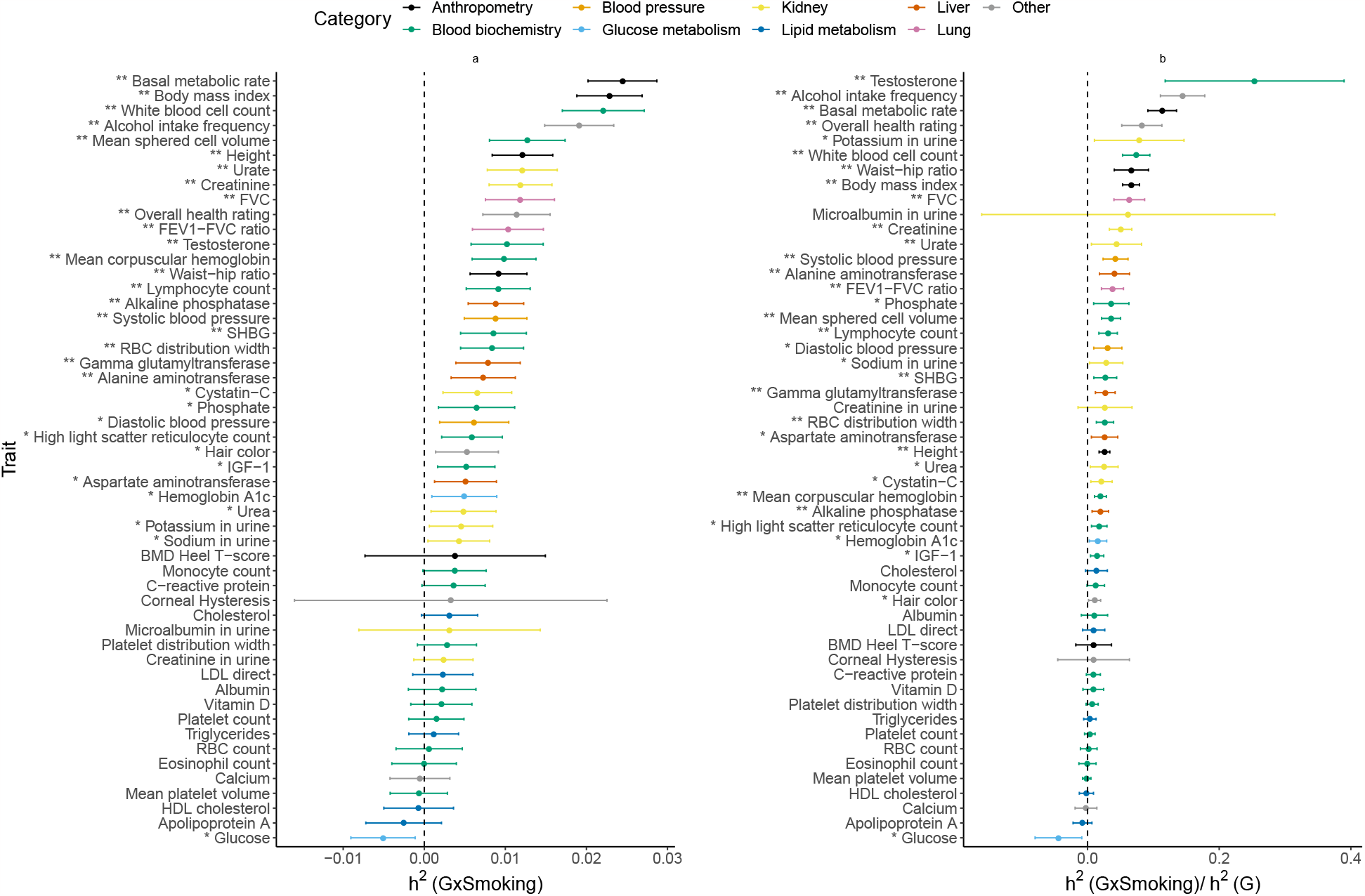
Estimates of GxSmoking heritability across phenotypes in UK Biobank. **a**) GxSmoking heritability and **b**) the ratio of GxSmoking to additive heritability. We applied GENIE to *N* = 291, 273 unrelated white British individuals and *M* = 459, 792 array SNPs (MAF*≥* 1%). Our model includes the environmental variable as a fixed effect and accounts for environmental heterogeneity. The environmental variable is standardized in these analyses. Error bars mark *±*2 standard errors centered on the point estimates. The asterisk and double asterisk correspond to the nominal *p <* 0.05 and *p* < 0.05/200, respectively.

#### Gene-by-Sex Interaction

We find 29 traits with statistically significant 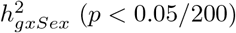 with 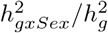 observed to be 8.7% on average (Figure 5). Serum testosterone levels showed the largest 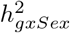 of 11% with the 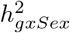 nearly as large as 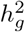 consistent with prior work showing differences in genetic associations [22, 23] and heritability [24] across males and females. Beyond testosterone, we observe significant 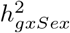 for several anthropometric traits, such as waist-hip-ratio adjusted for BMI (WHR) (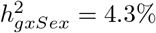and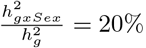), and lipid measures (results consistent for binary encoding; Supplementary Figure S14) consistent with previous work documenting sexspecific differences in the genetic architecture of anthropometric traits [25, 26, 27, 28, 29, 24]. Consistent with prior GWAS that identified genetic variants with sex-dependent effects [30, 31], our analyses of serum urate levels show substantial point estimates of 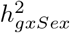, although these estimates are not statistically significant.

**Figure 5:**
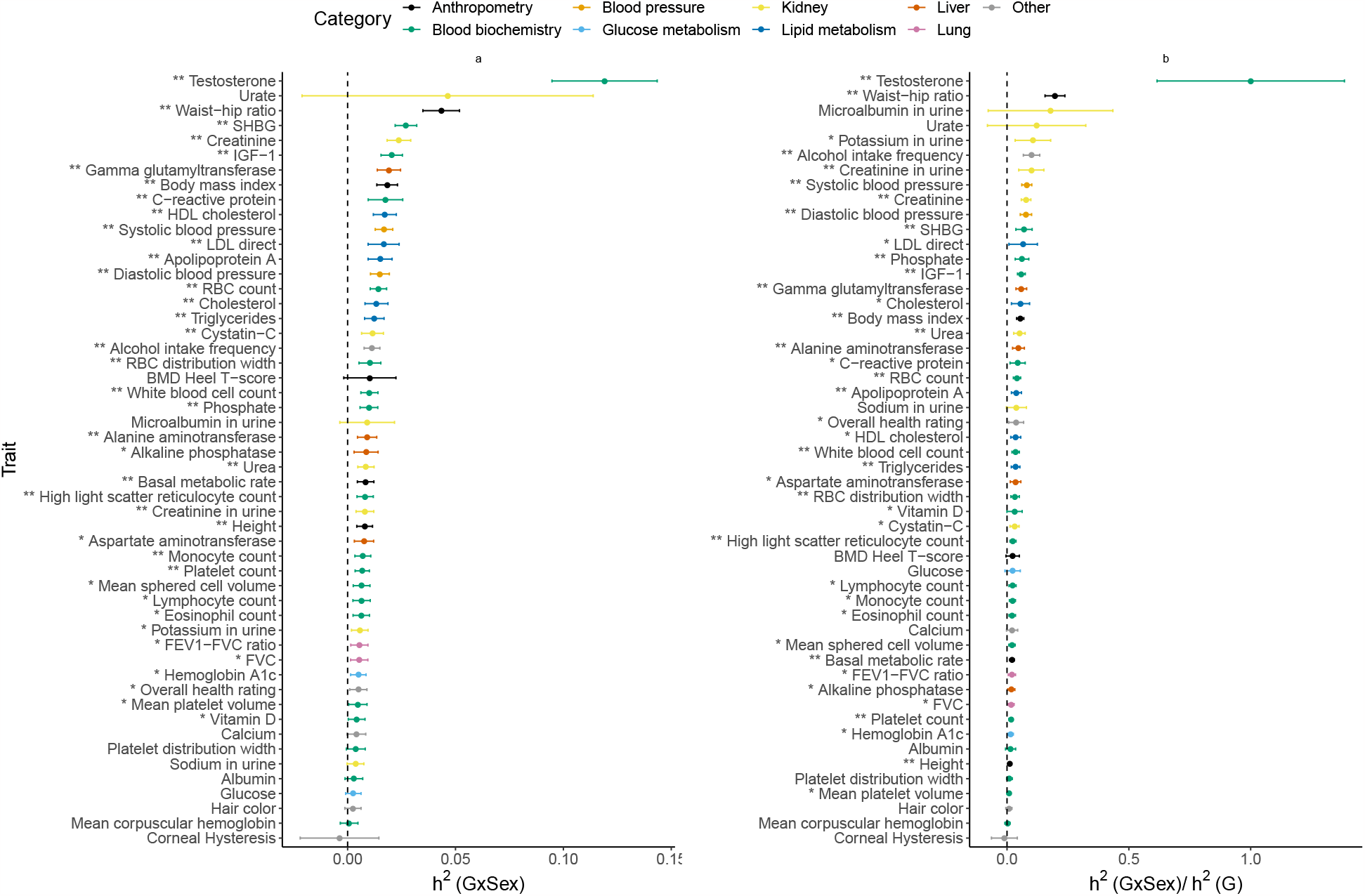
Estimates of GxSex heritability across phenotypes in UK Biobank. **a**) GxSex heritability and **b**) ratio of GxSex to additive heritability. We applied GENIE to *N* = 291, 273 unrelated white British individuals and *M* = 459, 792 array SNPs (MAF*≥* 1%). Our model includes the environmental variable as a fixed effect and accounts for environmental heterogeneity. The environmental variable is standardized in these analyses. Error bars mark *±*2 standard errors centered on the point estimates. The asterisk and double asterisk correspond to the nominal *p <* 0.05 and *p <* 0.05/200, respectively.

#### Gene-by-Age Interaction

We find 12 traits with statistically significant 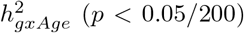 with 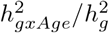 observed to be 4.3% on average (Figure 6). Lipid and blood pressure measures show some of the largest 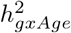 (about 2.5% for LDL-C and total cholesterol and 1.9% for diastolic blood pressure). Previous studies have found genetic variants in the SORT1 gene to have age-dependent effects on LDL cholesterol [32] and nominal evidence for age-dependent genetic effects on blood pressure regulation [33]. We find that BMI shows evidence for significant 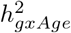 while WHR does not, expanding on prior work that identified age-dependent genetic variants for BMI but not for WHR in GWAS [26]. Interestingly, we used a standardized encoding of age so that GxAge effects capture the interaction of genetic effects on the phenotype as a function of deviation from the mean age in UKBB while previous studies typically focus on changes in genetic effects in bins of age. It is plausible that other codings of age, *e*.*g*., coding age to measure interactions as a function of older vs. younger individuals, could yield differing results.

**Figure 6:**
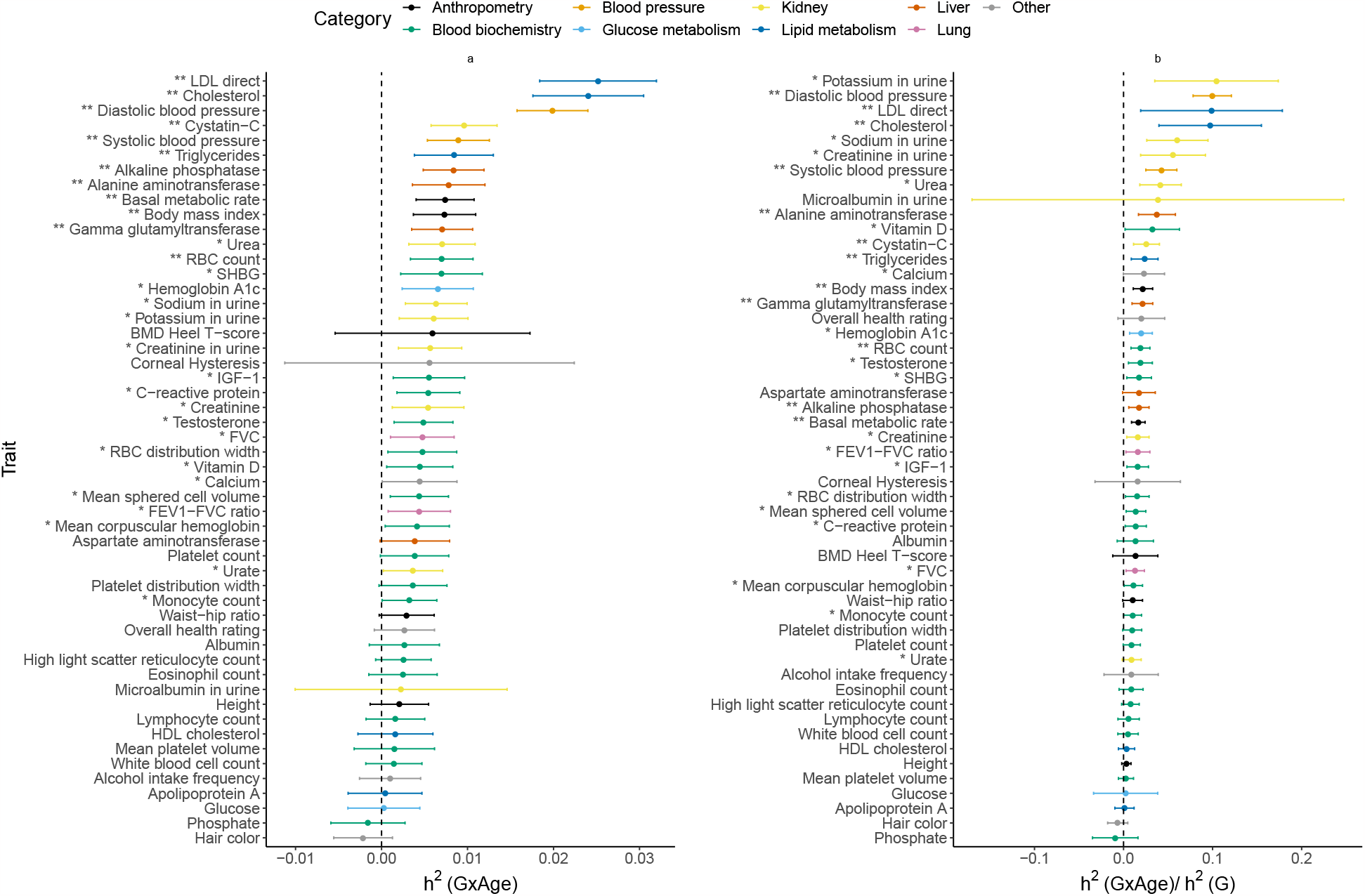
Estimates of GxAge heritability across phenotypes in UK Biobank. **a**) GxAge heritability and **b**) ratio of GxAge to additive heritability. We applied GENIE to *N* = 291, 273 unrelated white British individuals and *M* = 459, 792 array SNPs (MAF*≥* 1%). Our model includes the environmental variable as a fixed effect and accounts for environmental heterogeneity. The environmental variable is standardized in these analyses. Error bars mark *±*2 standard errors centered on the point estimates. The asterisk and double asterisk correspond to the nominal *p <* 0.05 and *p <* 0.05/200, respectively.

#### Gene-by-Statin Interaction

We find seven traits that show statistically significant evidence for 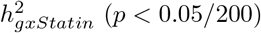 (*p <* 0.05*/*200) with an average ratio of 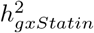 to 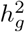 across traits of 5.2% (Figure 7). We find that LDL and total cholesterol show significant 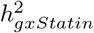 (1.7% and 1.6% respectively) while HDL cholesterol with a point estimate of 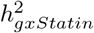 of 0.4% does not (results consistent for binary encoding; Supplementary Figure S15). We observe the largest estimates of 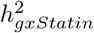 for HbA1c and blood glucose measurements (2% and 1.2% respectively) which are interesting in light of statin usage being shown to be associated with a small increase in risk for Type-2 Diabetes [34].

**Figure 7:**
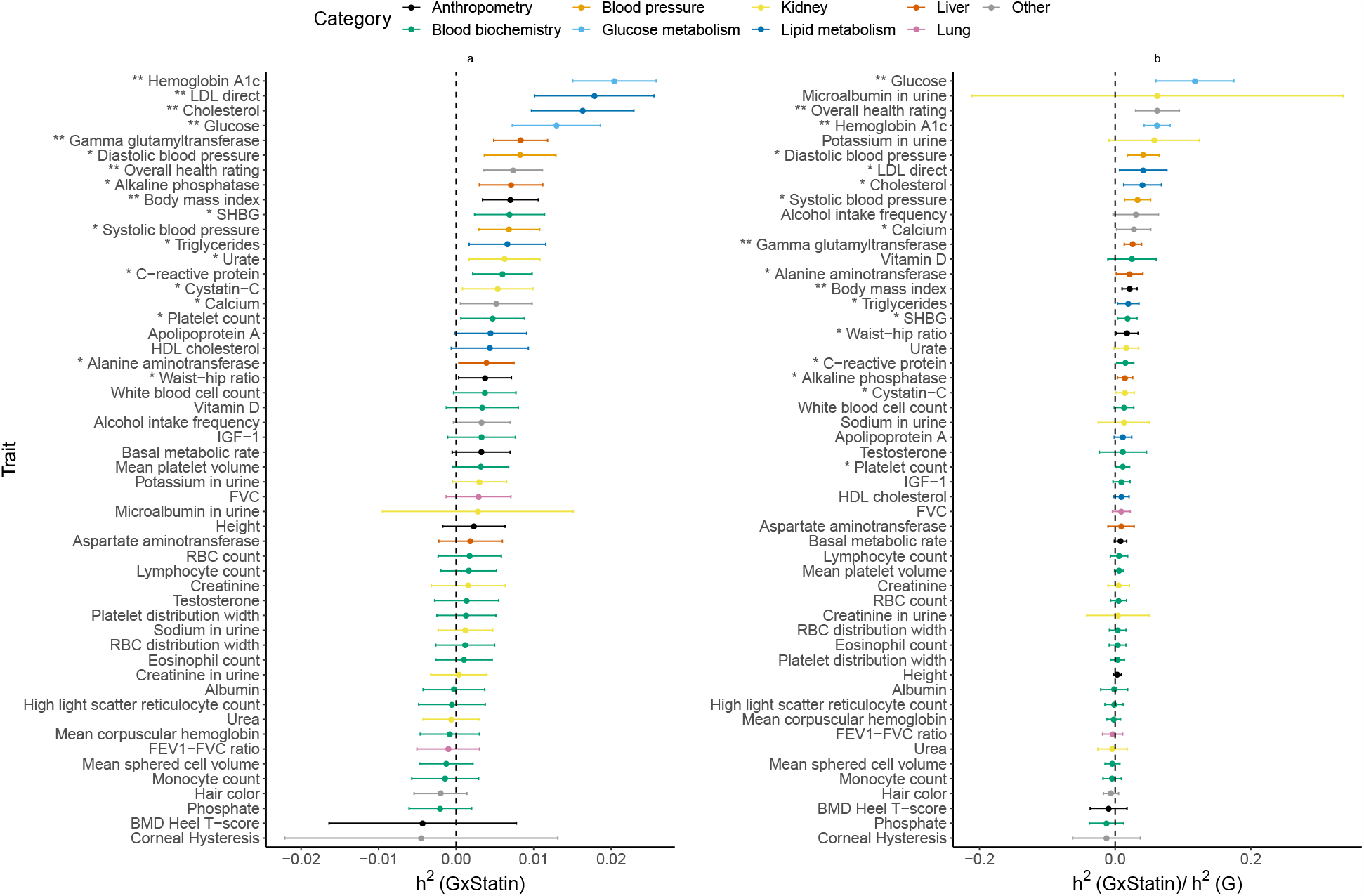
Estimates of GxStatin heritability across phenotypes in UK Biobank. **a**) GxStatin heritability and **b**) ratio of GxStatin to additive heritability. We applied GENIE to *N* = 291, 273 unrelated white British individuals and *M* = 459, 792 array SNPs (MAF*≥* 1%). Our model includes the environmental variable as a fixed effect and accounts for environmental heterogeneity. The environmental variable is standardized in these analyses. Error bars mark *±*2 standard errors centered on the point estimates. The asterisk and double asterisk correspond to the nominal *p <* 0.05 and *p <* 0.05/200, respectively.

#### Estimating GxE heritability from imputed SNPs

We applied GENIE to estimate 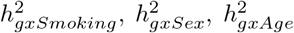,and 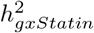 attributable to *M* = 7, 774, 235 imputed SNPs with MAF *≥* 0.1%. Prior work has shown that analyzing common and low-frequency variants with a single variance component can result in biased estimates of additive heritability [35, 36]. A solution to this problem involves fitting multiple variance components obtained by partitioning SNPs based on their frequency and local LD scores (as quantified by the LD-scores [37] or the LDAK scores [35]) [38, 36, 39, 40]. We follow this approach by partitioning SNPs into eight annotations based on quartiles of the LD-scores and two MAF annotations (MAF*<* 5% and MAF*>* 5%; Methods).

We performed simulations to show that GENIE applied with SNPs partitioned based on MAF and LD scores can accurately estimate 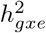 across varying MAF and LD-dependent genetic architectures while using a single component for all SNPs can lead to substantial biases (Supplementary Section S2; Supplementary Figure S16). We applied GENIE using MAF-LD partitions to jointly estimate 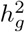 and 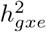 (Supplementary Figures S17, S18, S19, and S20). While estimates of 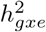 from imputed SNPs are largely concordant with the estimates obtained from array SNPs, we identify nine trait-E pairs for which the 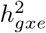 estimates are significantly different (*p <* 0.05*/*200). In all these cases, 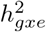 estimates from imputed SNPs are higher than those from array SNPs. For example, we estimate 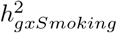 for BMI = 6.5 *±* 0.5% which is larger than our estimate based on array SNPs as well as a previous estimate of 4.0 *±* 0.8% based on common HapMap3 SNPs [5]. Restricting to traits with significant GxE in both array and imputed SNPs, we observed that the average ratio 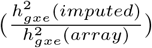 is 1.79 (2.31, 1.63, 1.17, and 1.17 respectively for GxSmoking, GxSex, GxAge, and GxStatin; Supplementary Figure S21). Across trait-E pairs with significant 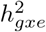, the average 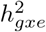 is 2.8% on the imputed data compared to 1.5% on array data while the ratio of 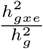 is 14.3% on the imputed data compared to 6.8% on the array data (averaged across trait-E pairs, we estimate 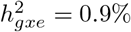 on imputed vs 0.7% on array data).

We explored the impact of fitting multiple variance components based on MAF and LD by applying GENIE to fit a single GxE and additive variance component using Smoking status as the environmental variable. While ten traits showed significant 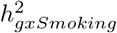 in both analyses, five traits were exclusively significant in the MAF-LD model while one was exclusively significant in the single-component model. Restricting to traits with significant GxSmoking in both models, 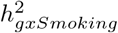 estimates in the MAF-LD model were about 3x those from the single-component model on average (Supplementary Figure S22). We also investigated whether MAF-LD partitioning affected estimates of 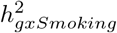 obtained from array SNPs. We find that 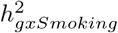 estimates are largely concordant whether obtained from a single component or a MAF-LD partitioned model (ratio of 0.99 on average) consistent with the array SNPs being relatively common (MAF*>* 1%).

Our analysis suggests that partitioning by MAF and LD is helpful for estimating 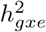 from both common and low-frequency SNPs and the inclusion low-frequency SNPs can increase estimates of 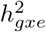 for specific traits.

#### Partitioning GxE heritability across MAF and LD annotations

Previous studies that have shown that the additive SNP effects increase with decreasing minor allele frequency (MAF) and local levels of linkage disequilibrium (LD) [41, 42, 43, 44], likely due to the effects of negative selection. However, the MAF-LD dependence of SNP effects in the context of specific environmental factors has not been empirically explored. Our analyses in the preceding section, showing differences in the genomewide 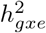 estimates when partitioning by MAF and LD vs. fitting a single variance component, suggest that GxE effects are expected to vary by MAF and LD in a pattern that is distinct from what would be expected when fitting a single variance component which assumes that the effect size at a SNP varies with its allele frequency *f* as 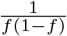 while not varying with local LD (for a fixed value of the allele frequency *f* ).

To explore the MAF-LD dependence of GxE effects, we used GENIE to partition 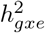 across MAF and LD annotations (while also simultaneously partitioning additive heritability) of *M* = 7, 774, 235 imputed SNPs divided into eight annotations based on quartiles of LD-scores and two MAF bins (low-frequency bins with MAF*<* 5% and high-frequency bins with MAF*≥* 5%). Within each of these eight bins, we defined the per-allele squared effect size as 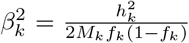 where 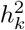 is the GxE (or additive) heritability attributed to bin *k, M*_*k*_ is the number of SNPs in bin *k* and *f*_*k*_ is the mean MAF in bin *k*.

For the sake of presentation, we selected one phenotype with high genome-wide GxE heritability for each of the four environmental variables analyzed (Figure 8; See the Supplementary Data 1 for results on all trait-E pairs). Across bins of MAF and LD, the magnitude of additive allelic effects tends to be larger than those of the GxE effects consistent with the genome-wide results. We observe that the per-allele squared GxE effect size 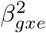 tends to increase with lower MAF within a given quartile of LD score and to increase with lower bins of LD score for a fixed MAF bin (Figure 8a). These trends are analogous to the relationship observed for additive per-allele effect sizes (Figure 8b). Across the trait-E pairs, restricting to the lowest quartile of LD scores, low-frequency SNPs tend to have higher per-allele GxE effect sizes compared to highfrequency SNPs: the ratio of 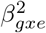 in low vs high MAF bins is 8.2 *±* 11.2, 24.6 *±* 19.7, 3.4 *±* 2.1, and 3.7 *±* 1.2 for HbA1c-statin, BMI-smoking, LDL-age, and testosterone-sex respectively. In the highest quartile of LD scores, we found no statistically significant differences in 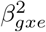 across low and high MAF SNPs in any of the four trait-E pairs (we also plot the per-standardized genotype additive and GxE heritability, 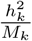, in Supplementary Figure S23).

**Figure 8:**
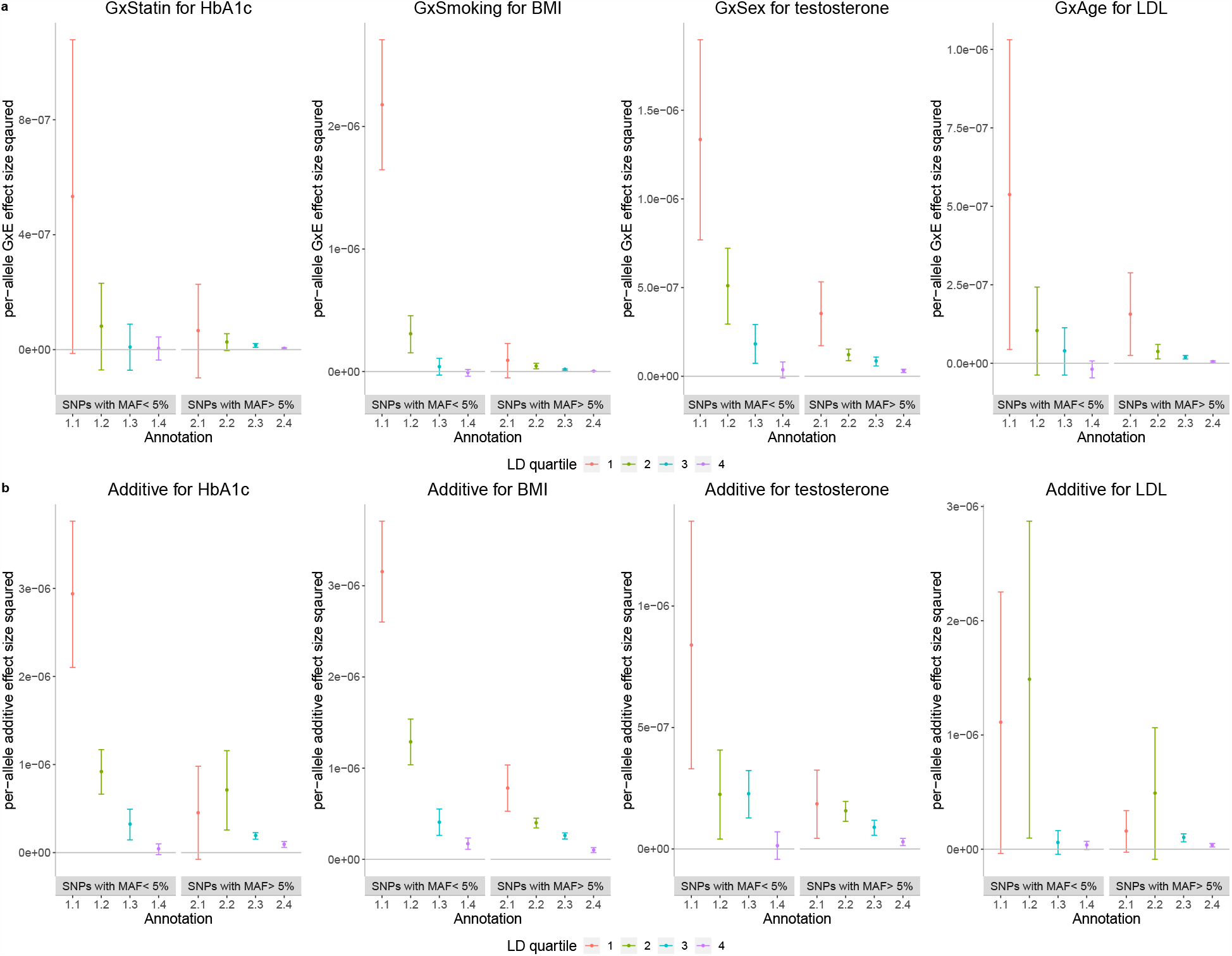
Per-allele squared GxE and additive effect sizes as a function of MAF and LD. **a**) The squared per-allele GxE effect size for four selected pairs of trait and environments (trait-E pairs). **b**) The squared per-allele additive effect size for the same trait-E pairs. The x-axis corresponds to MAF-LD annotations where annotation *i*.*j* includes SNPs in MAF bin *i* and LD quartile *j* where MAF bin 1 and MAF bin 2 correspond to SNPs with MAF *≤* 5% and MAF *>* 5% respectively while the first quartile of LD-scores correspond to SNPs with the lowest LD-scores respectively). The y-axis shows the per-allele GxE (or additive) effect size squared defined as 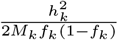 where 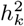 is the GxE (or additive) heritability attributed to bin *k, M*_*k*_ is the number of SNPs in bin *k*, and *f*_*k*_ is the mean MAF in bin *k*. Error bars mark *±*2 standard errors centered on the estimated effect sizes.

#### Partitioning GxE heritability across tissue-specific genes

The ability of GENIE to simultaneously estimate multiple, potentially overlapping, additive and GxE variance components enables us to explore how 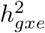 is localized across the genome. Specifically, we set to answer the question of whether 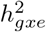 is enriched in genes specifically expressed in a given tissue as a means to identify tissues that are relevant to a trait in a specific environmental context.

We applied GENIE to estimate 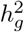 and 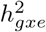 across each of 53 sets of genomic annotations defined as regions around genes that are highly expressed in a specific tissue in the GTEx dataset [17]. For each of the four environmental variables, we analyzed only traits with genome-wide significant 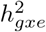 based on our prior analyses of the array SNPs. For every set of tissue-specific genes, we followed prior work [17] by jointly modeling the tissue-specific gene annotation as well as 28 genomic annotations that are part of the baseline LDSC annotations that include genic regions, enhancer regions, and conserved regions [45]). Specifically, our model has 29 additive variance components and 29 GxE variance components and estimates the additive and GxE heritability that can be attributed to genes specifically expressed in a tissue while controlling for the effects of the background annotations. A positive 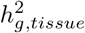 represents a positive contribution of genetic effects in a tissue to additive heritability [17]. Analogously, a positive 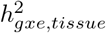 represents a positive contribution of genetic effects in this tissue to trait heritability in the context of the specific environment. We test estimates of 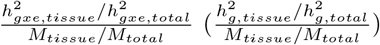 to answer whether a tissue of interest is enriched for GxE (additive) heritability conditional on the remaining genomic annotations included in the model (Methods). We first verified that our approach is able to detect previously reported enrichments for additive effects such as brain-specific enrichment for BMI and adipose-specific enrichment for WHR (Figure 9) [17]. Across 69 trait-E pairs with significant genome-wide GxE that we tested, we observed significant enrichment of 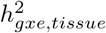 (FDR *<* 0.10) for at least one tissue in five trait-E pairs (we plot four of these pairs in Figure 9 since the results from the fifth LDL cholesterol-age are highly correlated with cholesterol-age). Across these trait-E pairs, we document differential patterns of enrichments for GxE effects compared to additive effects. BMI exhibits brain-specific enrichment of 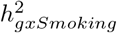 and 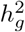 while WHR exhibits enrichment of 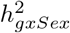 and 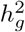 in adipose and breast tissue (in addition to the enrichment of 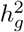 in the uterus and cardiovascular tissues). The adipose tissue-specific enrichment of 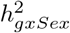 in WHR is notable in light of known instances of genes that are associated with WHR in adipose tissue in a sex-dependent manner. ADAMTS9, a gene involved in insulin sensitivity [25], is specifically expressed in adipose tissue and has been shown to be located near GWAS hits for WHR that are specific to females [46, 26, 25]. The transcription factor, KLF14, is located near a sex-dependent GWAS variant for WHR, type-2 Diabetes, and multiple other metabolic and anthropometric traits [47]. Further, the expression level of this gene is associated with the GWAS variant in adipose but not other tissues [47]. We also find instances where tissues that are enriched for 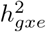 are distinct from those that are enriched for 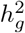. We observe that the enrichment of 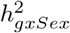 for basal metabolic rate in brain and adipose tissues is distinct from the tissues that are enriched in 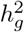 for the same trait (cardiovascular and digestive tissues) (Figure 9). Fitting this trend, we find that 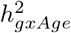 for cholesterol shows enrichment in cardiovascular tissues while 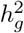 shows liver-specific enrichment. Finally, we find suggestive evidence that the liver is the most significantly enriched tissue for 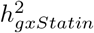 in HbA1c (*p* = 0.02) as well as for 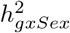 in testosterone (*p* = 0.005) although neither enrichment is significant at FDR of 0.10. These enrichments recapitulate known biology: the liver-specific enrichment of GxStatin effects for HbA1c reflect the tissues in which the target of statins (HMG-CoA-reductase) is expressed [48] while the liver-specific enrichment of GxSex for testosterone is consistent with previous findings implicating CYP3A7, a gene involved in testosterone metabolism that is specifically expressed in the liver and lies within a locus that contains one of the strongest GWAS signals for serum testosterone in females [23]

**Figure 9:**
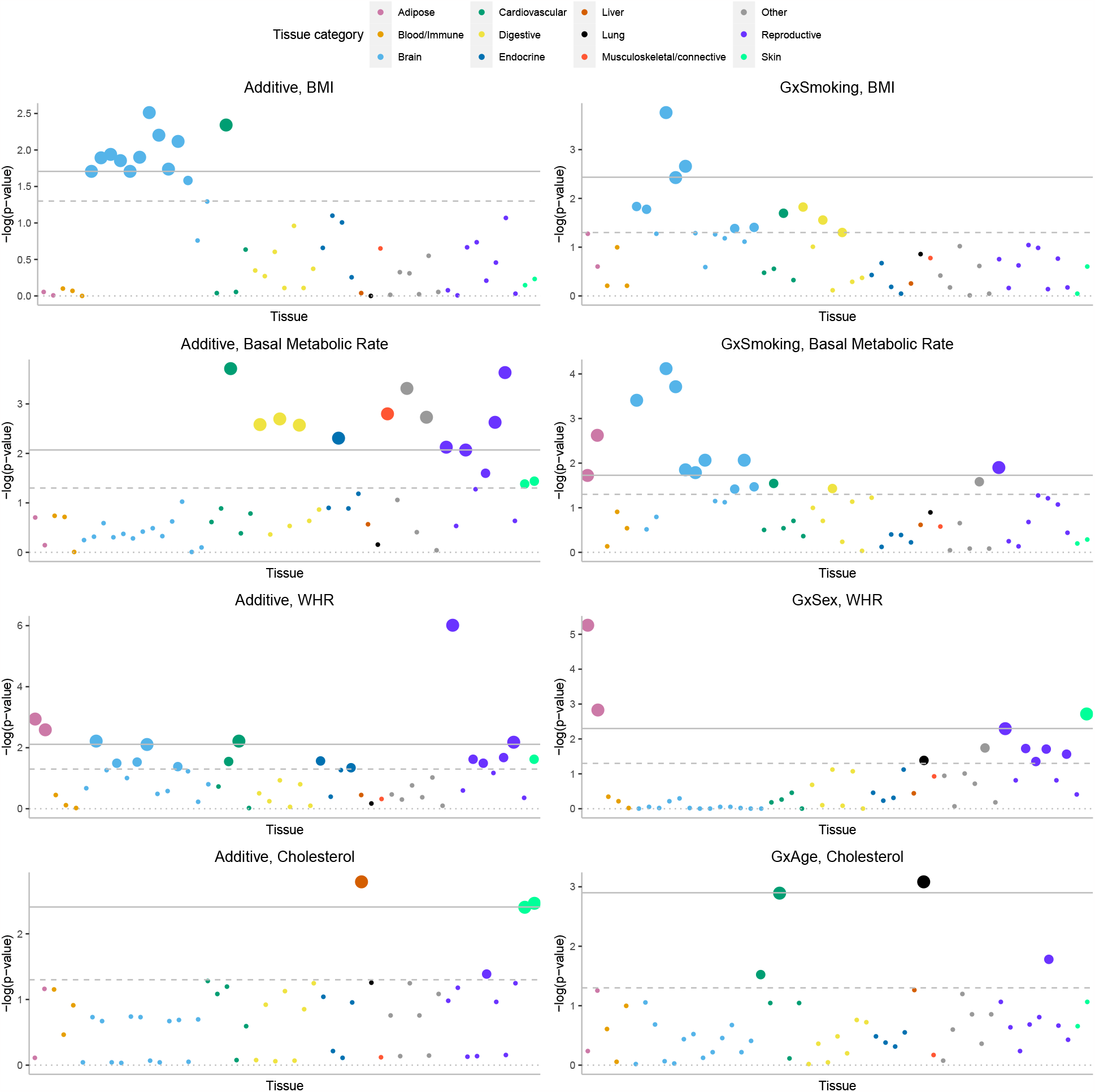
Partitioning GxE heritability across 53 tissue-specific genes.. We plot *−log*_10_(*p*) where *p* is the corresponding p-value of the tissue-specific GxE enrichment defined as 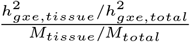. For every tissue specific annotation, we use GENIE to test whether this annotation is significantly enriched for per-SNP heritability, conditional on 28 functional annotations that are part of the baseline LDSC annotations. The dashed and solid lines correspond to the nominal *p <* 0.05 and FDR*<* 0.1 threshold, respectively.

## Discussion

We have described GENIE, a method that can jointly estimate the proportion of variation in a complex trait that can be attributed to GxE and additive genetic effects. GENIE can also partition GxE heritability across the genome with respect to annotations, such as functional and tissue-specific annotations or annotations defined based on the minor allele frequency (MAF) and local linkage disequilibrium (LD score) of each SNP to localize signals of GxE. GENIE provides well-calibrated tests for the existence of a GxE effect and has high power to detect GxE effects while being scalable to large datasets.

Our simulations and real data analysis results confirm the importance of including noise heterogeneity in GxE models. In UKBB data analyses, we observed about half of trait-E pairs with significant 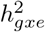 under the G+GxE model are no longer significant under the G+GxE+NxE model. Consistent with this observation, we estimate a substantial contribution of noise heterogeneity to trait variation.

After accounting for noise heterogeneity, we observe significant genome-wide 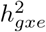 across more than a quarter of the trait-E pairs analyzed. Our finding has implications for understanding trait heritability by moving beyond the definition of narrow-sense heritability that only includes additive genetic effects. Based on our analyses, it is conceivable that approaches that can jointly model the hundreds of environmental variables measured in Biobank-scale datasets will further increase estimates of 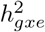. Additionally, our recovery of additional 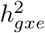 from low-frequency SNPs (0.1% *≥* MAF *<* 1%) point to traits where an understanding of GxE effects can benefit from whole-exome and whole-genome studies. Further, our results point to traits where GxE has the potential to improve genome-wide polygenic scores (GPS) of complex traits (since 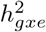 quantifies the maximum predictive accuracy that is achievable by a linear predictor based on GxE effects). In the context of sex as an environmental variable, sex-specific GPS has been shown to provide improved accuracy over agnostic scores [49, 50, 29, 24]. GxE has also been recently proposed as a possible explanation for why GPS may not generalize beyond the cohort on which these predictors were trained [6] so that modeling GxE in relevant traits could improve their transferability. Our finding that allelic effects for GxE increase with decreasing MAF and LD analogous to the relationship observed for additive allelic effects motivates an evolutionary understanding of these trends and can inform what we expect to learn from studies of rare genetic variation. Finally, our identification of sets of genes that are enriched for GxE can offer clues on trait-relevant tissues and pathways and has the potential to inform functional genomic studies [51, 52].

We discuss the limitations of our work as well as directions for future research. First, GENIE does not explicitly model G-E correlations [13]. While such correlations can lead to biases in estimates of GxE in the fixed-effect setting [53], it has been shown that, in the polygenic setting, the GxE variance component estimates remain unbiased when G-E correlations are independent of the polygenic GxE effects [14]. Nevertheless, there are plausible settings, where such correlations can lead to false positive or biased estimates of GxE, *e*.*g*., where the phenotype directly affects the environmental variable. Developing scalable methods that are accurate in these settings is an important direction for future work. Second, estimates of GxE heritability are sensitive to the scale on which traits and environmental variables are measured and how environmental variables are encoded. In this work, we analyze quantile-normalized traits (following prior studies) and encode discrete environmental variables using a univariate parameterization (either as a 0-1 vector for each environmental variable or as a standardized version). It might be preferable to work with traits measured on their original scale and to encode each level of discrete environmental variables by a separate 0-1 covariate (leading to *k* environmental covariates for a *k*-valued environmental variable). While such choices would necessarily be guided by domain knowledge and interpretability, GENIE supports easy-to-use and rapid exploration of the consequences of these choices and can aid in assessing the robustness of these choices (we have explored a limited space of these choices here). Third, the environmental variable relevant for GxE may not be measured directly or accurately so that the environmental variable that is measured in a dataset is best viewed as a proxy for the relevant latent environmental covariate. On a related note, while GENIE can model the impact of heterogeneous noise resulting from observed environmental variables by introducing NxE components, it is important to note that the heterogeneous noise may also arise due to non-observed environmental variables. Several recent works have tried to test for GxE when the environmental variables are not observed [10, 54]. These issues along with the possibility of reverse causality, *i*.*e*., where the trait affects the environmental variable, warrant caution in any causal interpretation of our results (although it might be possible to overcome some of these limitations in specific analyses such as GxSex). Fourth, the model underlying GENIE is not applicable to binary traits (either with or without ascertainment). GENIE can be extended to be applicable to binary traits (*e*.*g*., disease status) along the lines proposed in the context of additive [55, 56] and GxE estimation [14].

## Methods

### Generalized GxE linear mixed model

Let ***X*** denote a *N × M* genotype matrix, ***E*** denote a *N × L* matrix of environmental variables, ***C*** denote a *N × P* matrix of fixed-effect covariates, and ***y*** denote a *N* -vector of phenotypes. We assume the following linear mixed model:

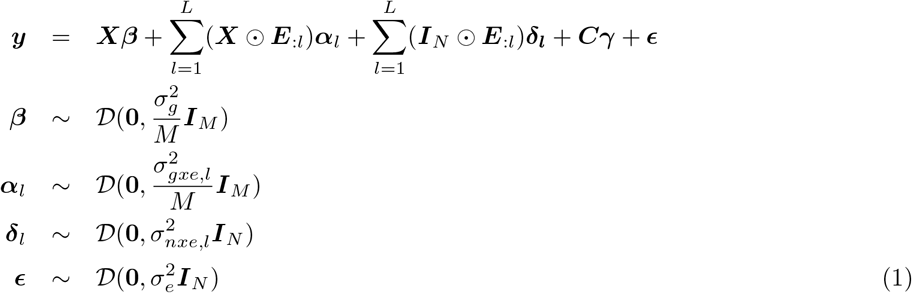

Here *D*(***µ*, Σ**) denotes an arbitrary distribution with mean ***µ*** and covariance **Σ, *E***_:*l*_ denotes *l*-th column of ***E***, and ⊙ denotes row-wise Kronecker product. ***β*** denotes the *M* -vector of SNP effect sizes, ***α***_*l*_ denotes the *M* -vector of genetic effect sizes in the context of environment *l* (GxE effects) while ***δ***_*l*_ denotes the *N* -vector of NxE effect sizes for environment *l*, and ***E*** denotes the *N* -vector of noise. 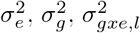, and 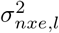 denote the residual variance, additive genetic, gene-by-environment, and noise-by-environment variance components respectively. These variance components can then be transformed into the additive heritability or the proportion of variance explained by additive effects (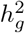 associated with 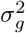) and the GxE heritability or the proportion of variance explained by interactions of genetics with a given environment (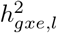 associated with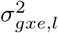).

### Estimation in the GxE linear mixed model

We assume without loss of generality that ***y*** is centered, and the columns of ***X*** and ***E*** are standardized. To estimate the variance components of our LMM, we use a Method-of-Moments (MoM) estimator that searches for parameter values so that the population moments are close to the sample moments. Since 𝔼 [***y***] = 0, we derived the MoM estimates by equating the population covariance to the empirical covariance. For simplicity, we exclude the matrix of covariates ***C*** from the model in the following derivation as the covariates can be efficiently projected out of the phenotype, genotypes, and interaction terms with minimal additional cost (Supplementary Section S1).

For compactness, we denote ***Z***_0_ = ***X, Z***_*l*_ = ***X ⊙ E***_***l***_ for *l* = 1, .., *L*, ***Z***_*l*_ = ***I***_***N***_ ***⊙ E***_**:*l***_ for *l* = *L* + 1, .., 2*L*, and ***Z***_2*L*+1_ = ***I***_*N*_ . The population covariance is given by:

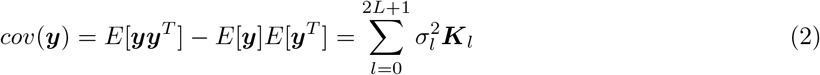

where

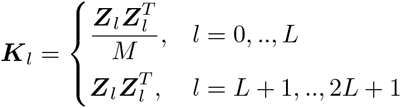

and

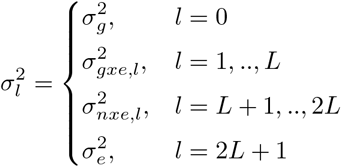

Using ***yy***^T^ as our estimate of the empirical covariance, we need to solve the following least squares problem to find the variance components.

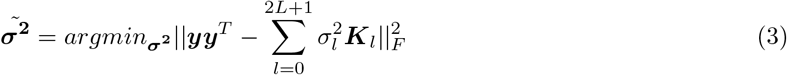

The MoM estimator satisfies the following normal equations:

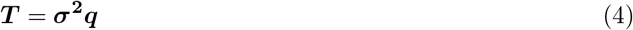

where ***T*** is matrix with entries *T*_*ij*_ = *tr*(***K***_*i*_***K***_*j*_), *i, j ∈ {*0, .., 2*L* + 1*}*, and ***q*** and 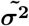 are vectors with entries

*c*_*l*_ = ***y***^*T*^ ***K***_*l*_***y***, for *l ∈ {*0, .., 2*L* + 1*}*.

The heritability associated with component *i* for a component that represents additive genetic or GxE effects (equivalently, the proportion of variance explained by component *i*) is defined as follows:

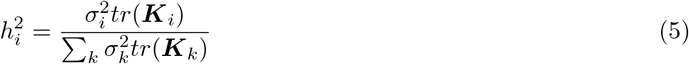

The aforementioned definition of heritability holds when the matrices ***Z***’s columns have zero means, and *N* is large. To explicitly ensure that the columns of GxE matrices also have zero means, a column consisting of all ones is included in the covariate matrix. Consequently, when the covariates are projected out of the GxE matrices (Supplementary Section S1), it guarantees that all columns have zero means.

### Computational challenges

Computing the coefficients of the system of linear equation 4 presents computational challenges. The main computational bottleneck is the evaluation of the quantities *T*_*ij*_ for *i, j ∈ {*0, …, 2*L* + 1*}* which requires *O*(*N* ^2^*M* ). Therefore, the total time complexity for exact MoM is *O*(*N* ^2^*ML* + *L*^3^) imposing challenging memory or computation requirements for Biobank-scale data (*N* in the hundreds of thousands, *M* in the millions, *L* in the hundreds or thousands).

### Scalable estimation

Instead of computing the exact value of *T*_*ij*_, GENIE uses a randomized estimator of the trace [57]. This estimator uses the fact that for a given *N × N* matrix ***C, w***^*T*^ ***Cw*** is an unbiased estimator of *tr*(***C***) (*E*[***w***^*T*^ ***Cw***] = *tr*[***C***]) where ***w*** be a random vector with mean zero and covariance ***I***_*N*_ . Hence, we can estimate the values *T*_*ij*_, *i, j ∈ {*0, …, 2*L* + 1*}* as follows:

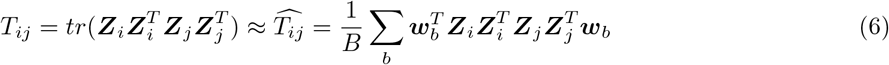

Here ***w***_1_, …, ***w***_*B*_ are *B* independent random vectors with zero mean and covariance ***I***_*N*_ . In GENIE, we draw these random vectors independently from a standard normal distribution. Note that computing *T*_*ij*_ by using the above estimator involves matrix-vector multiplications which are repeated *B* times. Therefore, the total running time is *O*(*LNMB*).

Moreover, we can leverage the structure of the genotype matrix which only contains entries in *{*0, 1, 2*}*. For a fixed genotype matrix ***X***_*k*_, we can improve the per iteration time complexity of matrix-vector multiplication from *𝒪* (*NM* ) to 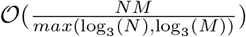 by using the Mailman algorithm [58]. Solving the normal equations takes 𝒪 (*L*^3^) time so that for a small number of components (*L*), the overall time complexity of our algorithm is 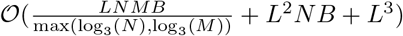.

#### Standard errors of the estimates and inclusion of covariates

We proposed two ways to compute standard errors of the estimates. First, we can use a computationally efficient block Jackknife [42], which does not require any assumptions on the distribution of the effect sizes. We also propose an efficient approach to compute standard errors based on the assumption that the phenotype is normally distributed (see Supplementary Notes). In our real data analysis, we used the block Jackknife to compute the standard errors of the estimates with 100 blocks defined over SNPs.

##### Partitioning GxE heritability across the genome

Although the model defined in Equation 1 is beneficial in quantifying the total GxE effects for a given E, it is interesting to identify and interpret the interaction of E with specific regions of the genome, such as SNPs with a particular range of minor allele frequencies or SNPs that lie within genes expressed specifically in a tissue. Following our previous work [42], the genotype component **X** can be assigned to T (potentially overlapping) components with respect to a set of annotations (such as MAF/LD or functional annotations). Thus, we extend our model as follows:

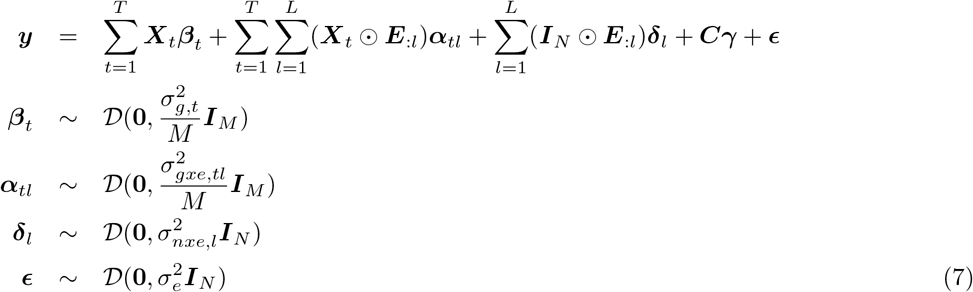

Here ***X***_*t*_ is the genotype of annotation *t* with *M*_*t*_ SNPs, ***α***_*tl*_ refers to the effect sizes of SNPs in annotation *t* in the context of environment *l*. Analogously, 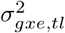 refers to the variance component for SNPs in annotation *t* in the context of environment *l* while 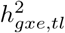 refers to the GxE heritability associated with annotation *t* in the context of environment *l*.

Given estimated GxE heritabilties under the above model, we define the enrichment of genetic effects in annotation *t* in the context of environment *l* (also termed GxE enrichment) as follows :

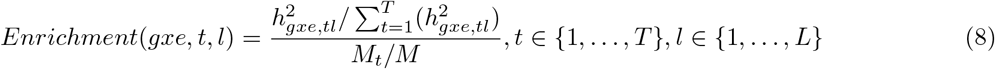

GENIE uses the same randomized trace estimator approach to efficiently estimate GxE heritability.

### Estimating GxE in the UK Biobank

We applied GENIE to the UK Biobank (UKBB) [8] where we considered environmental variables such as smoking status, sex, and statin medication. For every environmental variable, we applied GENIE to estimate additive heritability 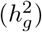 and GxE heritability 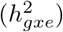 across 50 quantitative phenotypes (in a model that included the environmental variable as a main effect and accounted for environmental heterogeneity). In this study, we restricted our analysis to SNPs that were present in the UK Biobank Axiom array used to genotype the UK Biobank. SNPs with greater than 1% missingness and minor allele frequency smaller than 1% were removed. Moreover, SNPs that failed the Hardy-Weinberg test at significance threshold 10^*−*7^ were removed. We restricted our study to self-reported British white ancestry individuals who are *>* 3^*rd*^ degree relatives that are defined as pairs of individuals with kinship coefficient *<* 1*/*2^(9*/*2)^ [8]. Furthermore, we removed individuals who are outliers for genotype heterozygosity and/or missingness. Finally, we obtained a set of *N* = 291, 273 individuals and *M* = 459, 792 SNPs to use in the real data analyses.

We included age, sex, age^2^,age*×*sex, age^2^*×*sex, and the top 20 genetic principal components (PCs) as covariates in our analysis for all traits. We always include the environmental variable as a covariate in these analyses. We used PCs precomputed by the UK Biobank from a superset of 488, 295 individuals. Additional covariates were used for waist-to-hip ratio (adjusted for BMI) and diastolic/systolic blood pressure (adjusted for cholesterol-lowering medication, blood pressure medication, insulin, hormone replacement therapy, and oral contraceptives). We standardized environmental variables in our primary analyses. The standardized coding for binary environmental variables has an invariant property in the sense that the covariance matrix would be the same regardless of flipping the 0*/*1 coding. We also considered binary coding of environmental variables was relevant. Statin usage is defined as a binary environmental variable based on C10AA (the American Therapeutic Chemical (ATC) code of statin), which corresponds to taking any subtype of statin medications. Smoking status is defined as a categorical variable with three possible values (never, previous, current).

We considered an additional analysis of genotypes at high-quality imputed SNPs (with a hard call threshold of 0.2 and an INFO score *≥* 0.8) with MAF*≥* 0.1% in the *N* = 291, 273 unrelated white British individuals. We further restricted our analyses to SNPs that are under Hardy-Weinberg equilibrium (*p <* 10^*−*7^) and are confidently imputed in more than 99% of the individuals. Additionally, we excluded SNPs in the MHC region, resulting in a total of *M* = 7, 774, 235 SNPs.

In our analysis of heritability partitioned based on MAF-LD annotations, we divided SNPs into eight annotations based on quartiles of the LD scores (computed in-sample using GCTA) and two MAF bins (MAF*<* 5% and MAF*≥* 5%). In our analyses of heritability partitioned based on tissue-specific gene expression annotations, we used the annotations for the 53 tissue-specific genes generated by Finucane et al. [17] using a matrix of normalized gene expression values from the Genotype-Tissue Expression (GTEx) database, which included samples from various tissues, including the focal tissue. The authors calculated a t-statistic for each gene to determine its specific expression in the focal tissue and ranked all genes based on their t-statistic. They defined the top 10% of genes with the highest t-statistic as the set of specifically expressed genes for the focal tissue. To improve the accuracy of the gene set construction, 100-kb windows are added on either side of the transcribed region of each gene in the set of specifically expressed genes to generate a genome annotation that corresponds to the focal tissue.

## Data Availability

Access to the UK Biobank resource is available via application at: http://www.ukbiobank.ac.uk.

## Code Availability

GENIE software is an open-source software freely available at https://github.com/sriramlab/GENIE.

## Acknowledgments

This research was conducted using the UK Biobank Resource under application 331277. We thank the participants of UK Biobank for making this work possible. This work was funded by NIH grants R35GM125055 (A.P. and S.S.), HG006399 (S.S.), and NSF grant CAREER-1943497 (A.P. and S.S.).

The authors would like to thank Alkes Price and Arbel Harpak for their feedback on the manuscript. The authors would also like to acknowledge the stimulating discussions at the UCLA Computational Genomics Summer Institutes (supported by NIH grants GM135043 and GM112625) and the 2018 Bertinoro workshop in Statistical and Computational Genomics that enabled this work.

