## Supplementary materials for "A scalable and robust variance components method reveals insights into the architecture of gene-environment interactions underlying complex traits"

### Supplementary Notes

#### S1 Including covariates

We can extend each of our models to include covariates as follows:

$$\boldsymbol{y} = \boldsymbol{W}\boldsymbol{\alpha} + \sum_k \boldsymbol{Z}_k\boldsymbol{\beta}_k + \boldsymbol{\epsilon} \tag{9}$$

Here $\boldsymbol{W}$ is a $N \times C$ matrix of covariates while $\boldsymbol{\alpha}$ is a vector of fixed effects of length $C$. Matrix $\boldsymbol{Z}_k$ can represent genotype or GxE matrices . In this setting, we need to solve the following normal equations to estimate the variance components.

$$\begin{bmatrix} \boldsymbol{T} & \boldsymbol{b} \\ \boldsymbol{b}^T & N - C \end{bmatrix} \begin{bmatrix} \sigma_1^2 \\ \vdots \\ \sigma_k^2 \\ \sigma_e^2 \end{bmatrix} = \begin{bmatrix} \boldsymbol{c} \\ \boldsymbol{y}^T\boldsymbol{V}\boldsymbol{y} \end{bmatrix} \tag{10}$$

Here $\boldsymbol{V} = \boldsymbol{I}_N - \boldsymbol{W}(\boldsymbol{W}^T\boldsymbol{W})^{-1}\boldsymbol{W}^T$ and $\boldsymbol{T}$ is a $K \times K$ matrix where $T_{k,l} = tr(\boldsymbol{K}_k\boldsymbol{V}\boldsymbol{K}_l\boldsymbol{V})$, and $b$ is a vector of length $K$ where $b_k = tr(\boldsymbol{V}\boldsymbol{K}_k)$, and $\boldsymbol{c}$ is a vector of length $K$ where $c_k = \boldsymbol{y}^T\boldsymbol{V}\boldsymbol{K}_k\boldsymbol{V}\boldsymbol{y}$, $\boldsymbol{K}_k = \frac{\boldsymbol{Z}_k\boldsymbol{Z}_k^T}{M}$ where $M$ is the number of column in $Z_k$. Commonly, the number of covariates $C$ is small (tens to hundreds) so including covariates does not significantly affect the computational cost. The cost of computing the elements of the normal equations 10 includes the cost of inverting $\boldsymbol{W}^T\boldsymbol{W}$ which is a $C \times C$ matrix and multiplying $\boldsymbol{W}$ by a real-valued vector of length $N$ can be computed in $\mathcal{O}(C^3 + NC)$.

#### S2 Simulations of MAF and LD-dependent genomic architectures

To simulate MAF and LD-dependent architectures, we simulated phenotypes from genotypes using the following model that extends prior models of additive genetic architecture [36, 59] to include GxE effects:

$$\begin{aligned} \sigma_{g,m}^2 &= Sc_m w_m^b [f_m(1-f_m)]^a \\ (\boldsymbol{\beta}_1, \boldsymbol{\beta}_2, .., \boldsymbol{\beta}_m)^T &\sim \mathcal{N}(\boldsymbol{0}, diag(\sigma_{g,1}^2, \sigma_{g,2}^2, ..., \sigma_{g,m}^2)) \\ \sigma_{gxe,m}^2 &= S' c_m' w_m^b [f_m(1-f_m)]^a \\ (\boldsymbol{\alpha}_1, \boldsymbol{\alpha}_2, .., \boldsymbol{\alpha}_m)^T &\sim \mathcal{N}(\boldsymbol{0}, diag(\sigma_{gxe,1}^2, \sigma_{gxe,2}^2, ..., \sigma_{gxe,m}^2)) \\ \boldsymbol{y}|\boldsymbol{\beta}, \boldsymbol{\alpha} &\sim \mathcal{N}(\boldsymbol{X}\boldsymbol{\beta} + (\boldsymbol{X} \odot \boldsymbol{E})\boldsymbol{\alpha}, (1 - h_g^2 - h_{gxe}^2 - h_{nxe}^2)\boldsymbol{I}_N + h_{nxe}^2(\boldsymbol{I}_N \odot \boldsymbol{E})) \end{aligned} \tag{11}$$

where $h_g^2, h_{gxe}^2, h_{nxe}^2 \in [0, 1]$, $a \in \{0, 0.75\}$, $b \in \{0, 1\}$. Here $S$ and $S'$ are normalizing constants chosen so that $\sum_{m=1}^M \sigma_{g,m}^2 = h_g^2$, $\sum_{m=1}^M \sigma_{gxe,m}^2 = h_{gxe}^2$. Additive and GxE effect sizes are denoted by $\boldsymbol{\beta}$ and $\boldsymbol{\alpha}$ respectively. $f_m$ and $w_m$ are the minor allele frequency and LD score of $m^{th}$ SNP respectively. In this model, $c_m, c_m' \in \{0, 1\}$ are indicator variables for the causal status of SNP $m$ ($c_m = 1$ and $c_m' = 1$ for all SNPs). The LD score of a SNP is defined to be the sum of the squared correlation of the SNP with all other SNPs that lie within a specific distance. Setting $a = 0, b = 0$ for additive effects results in the GCTA model where the per-standardized genotypic effect sizes at a SNP do not vary with MAF or LD score [11, 37] while setting $a = 0.75, b = 1$ results in the LDAK model [35]. Our simulations assume that the GxE effects follow the same coupling as the additive effects. We used phenotypes from $N = 40k$ individuals genotyped at $M = 459,792$ SNPs on the UKBB array.

### Supplementary Tables

| Method | Variance component | Mean | SE | Bias | Test of bias p-value |
|---|---|---|---|---|---|
| GENIE | G | 0.2005 | 0.0382 | 5e-04 | 0.9006 |
| GENIE | GxE1 | -0.0021 | 0.027 | -0.0021 | 0.4302 |
| GENIE | GxE2 | 0.0035 | 0.0257 | 0.0035 | 0.1778 |
| GENIE | GxE3 | -1e-04 | 0.0275 | -1e-04 | 0.9692 |
| GENIE | GxE4 | 0.0016 | 0.0245 | 0.0016 | 0.5094 |
| GENIE | GxE5 | 0.0017 | 0.0267 | 0.0017 | 0.5159 |
| GENIE | GxE6 | 0.0974 | 0.0278 | -0.0026 | 0.3449 |
| GENIE | GxE7 | 0.0981 | 0.0266 | -0.0019 | 0.4673 |
| GENIE | GxE8 | 0.0975 | 0.0257 | -0.0025 | 0.3329 |
| GENIE | GxE9 | 0.0098 | 0.0263 | -2e-04 | 0.9537 |
| GENIE | GxE10 | 0.0094 | 0.0257 | -6e-04 | 0.8082 |

Table S1: **Accuracy of GENIE in the setting of multiple environmental variables**: We reported the bias, and SE of GENIE under different settings with $L = 10$ environmental variables. Bias, mean and SE are computed from 100 replicates. We report p-value of a test of the null hypothesis of no bias in the estimates of variance components.

#### Supplementary Figures

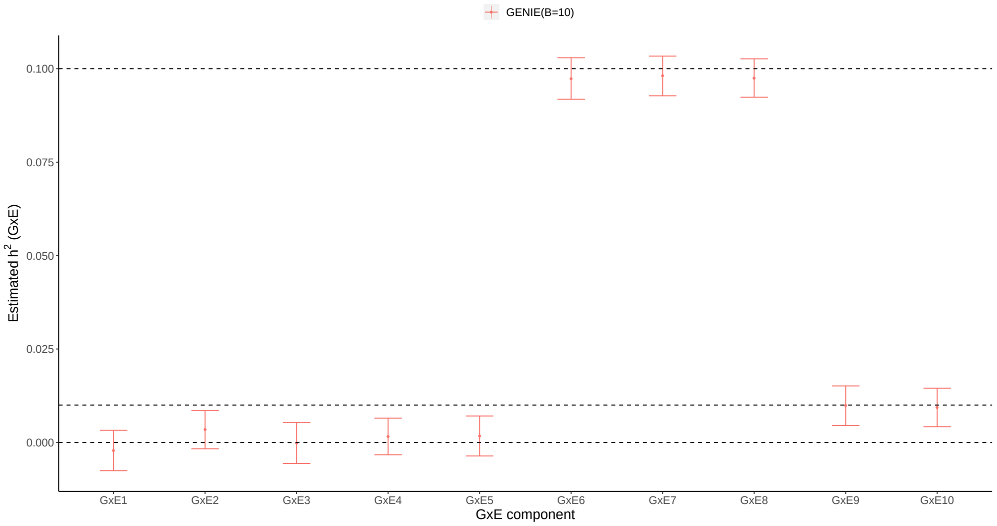


Figure S1: **Accuracy of GENIE when applied to multiple environmental variables**. We evaluated $h^2_{gxe}$ estimates of GENIE for different values of $\sigma^2_{gxe}$. We simulated phenotypes with 10 environmental variables where $\sigma^2_g = 0.2$, $\sigma^2_{ge1} = \sigma^2_{ge2} = \sigma^2_{ge3} = \sigma^2_{ge4} = \sigma^2_{ge5} = 0$, $\sigma^2_{ge6} = \sigma^2_{ge7} = \sigma^2_{ge8} = 0.10$ and $\sigma^2_{ge9} = \sigma^2_{ge10} = 0.01$. Points and error bars represent the mean and $\pm 2$ SE. Mean and SE are computed from 100 replicates. Here $B$ is the number of random vectors used by GENIE with $B = 10$ the value that we use as default (we reported values of means, SEs, and p-values of a test of the null hypothesis of no bias in the estimates of variance components in Supplementary Table 1)

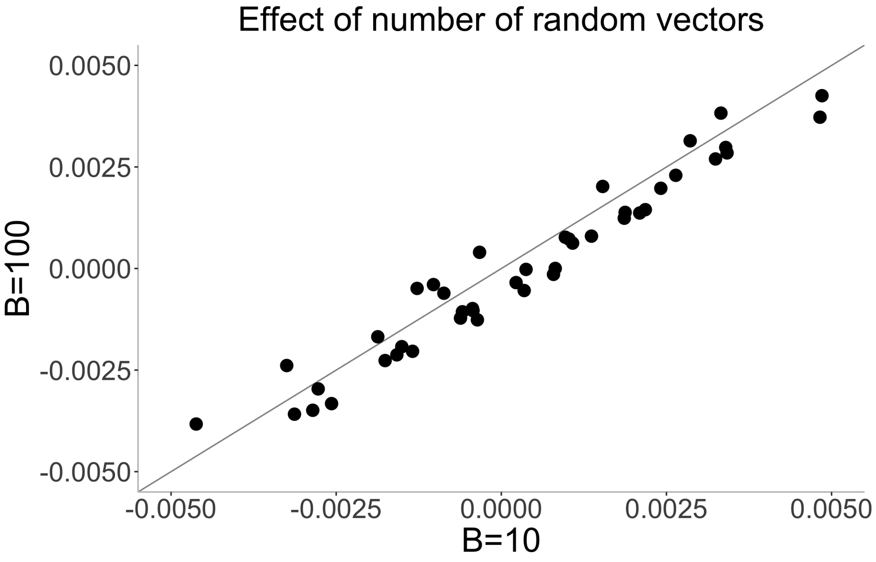


Figure S2: **Comparison of $h^2_{gxe}$ estimates with $B = 10$ and $B = 100$.** We simulated phenotypes from $M = 459,792$ array SNPs and $N = 291,273$ individuals where $h^2_g = 0.25$, $h^2_{gxe} = 0$ and the causal ratio is 10% .

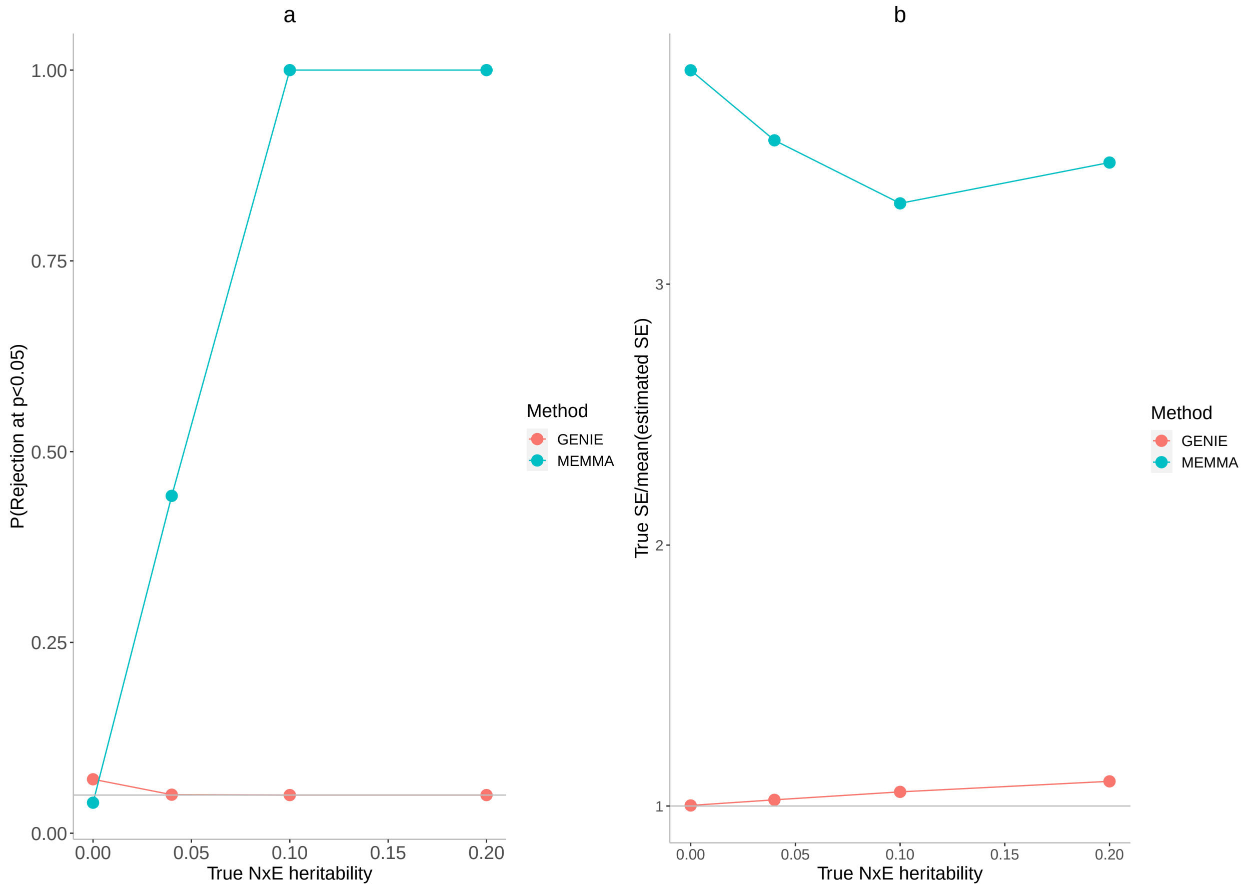


Figure S3: **Effect of estimated standard error on controlling false positive rate** . **a**) We assessed the calibration of GENIE and MEMMA using their true SE instead of the estimation of SE in simulations. MEMMA has biased estimates of SE, leading to a high false positive rate even in the absence of a NxE effect. **b**) We plot the ratio of true SE over the mean of estimated SE across 100 replicates as a function of the variance of the NxE effect.

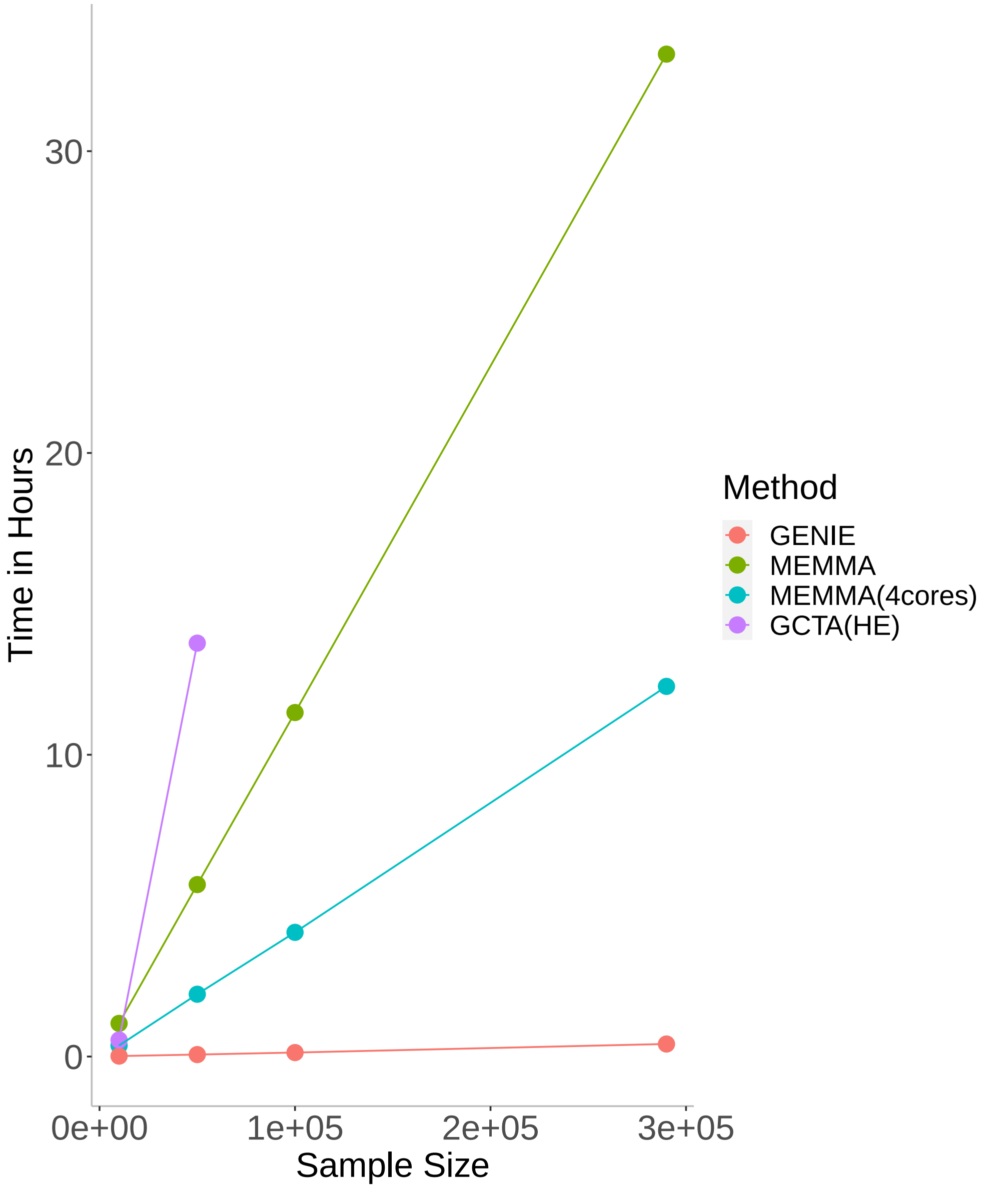


Figure S4: **Comparison of runtimes**. We evaluated the runtime of GENIE, MEMMA, and GCTA(HE) with increasing sample size $N$ (for a fixed number of SNPs $M = 459,792$ and single environmental variable). We fit single G and GxE variance components. All methods were run on an Intel(R) Xeon(R) Gold 6140 CPU 2.30GHz with 187 GB RAM. Ten random vectors are used by GENIE and MEMMA. The runtime of GCTA(HE) includes the computation of the GRM. GENIE and GCTA(HE) are executed on a single core while MEMMA is run on both a single core and four cores.

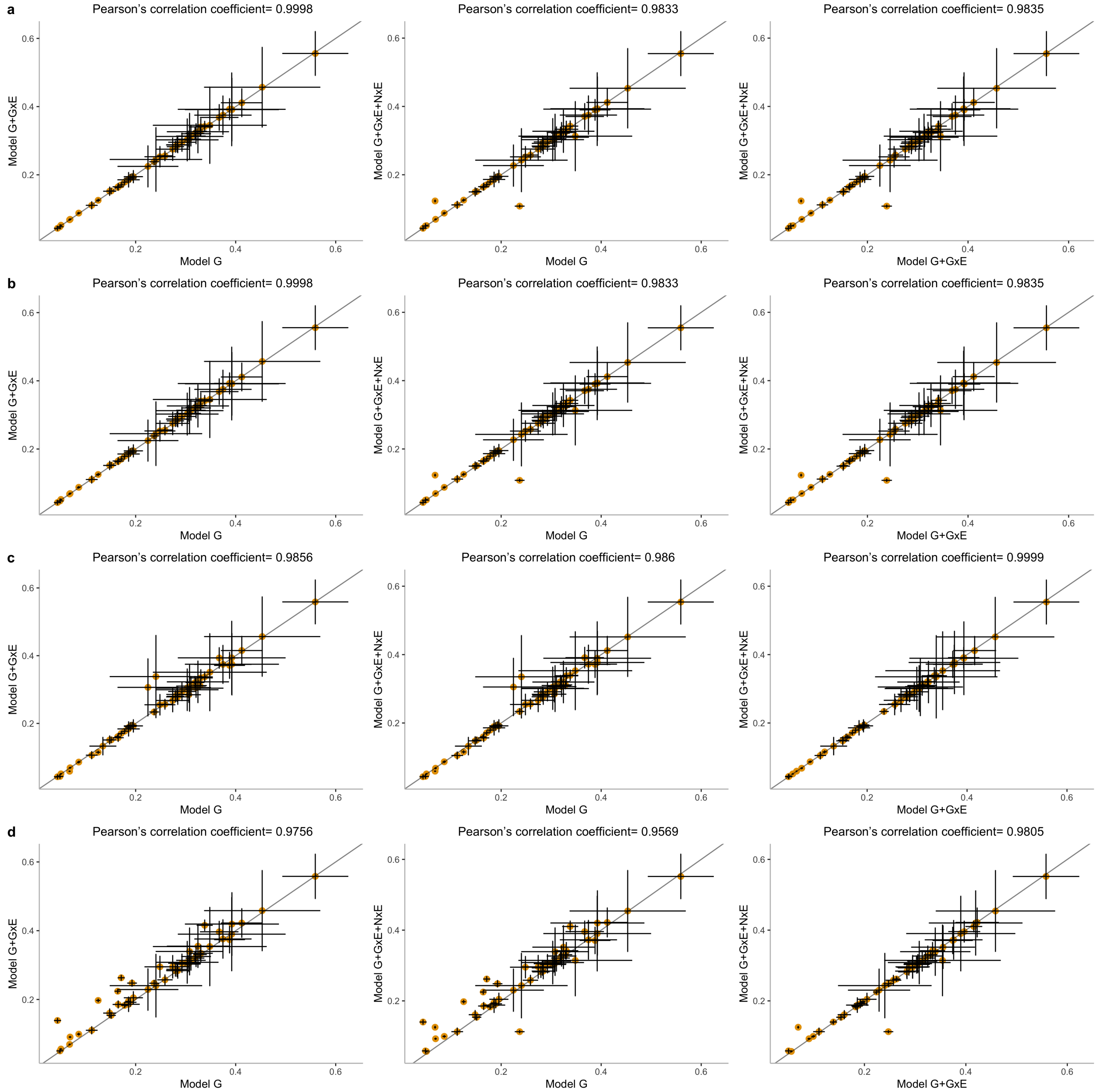


Figure S5: **Estimated additive heritability from three different models.** In this figure, G, GxE and NxE refer to additive, gene-by-environment, and noise-by-environment components, respectively. Every model is named by a set of variance components fitted jointly under that model. Estimates of additive components under the three models where the environmental variable is **a**) smoking status, **b**) sex, **c**) statin usage, and **d**) age. The estimates of additive heritability obtained by GENIE are consistent under these three models across environmental variables.

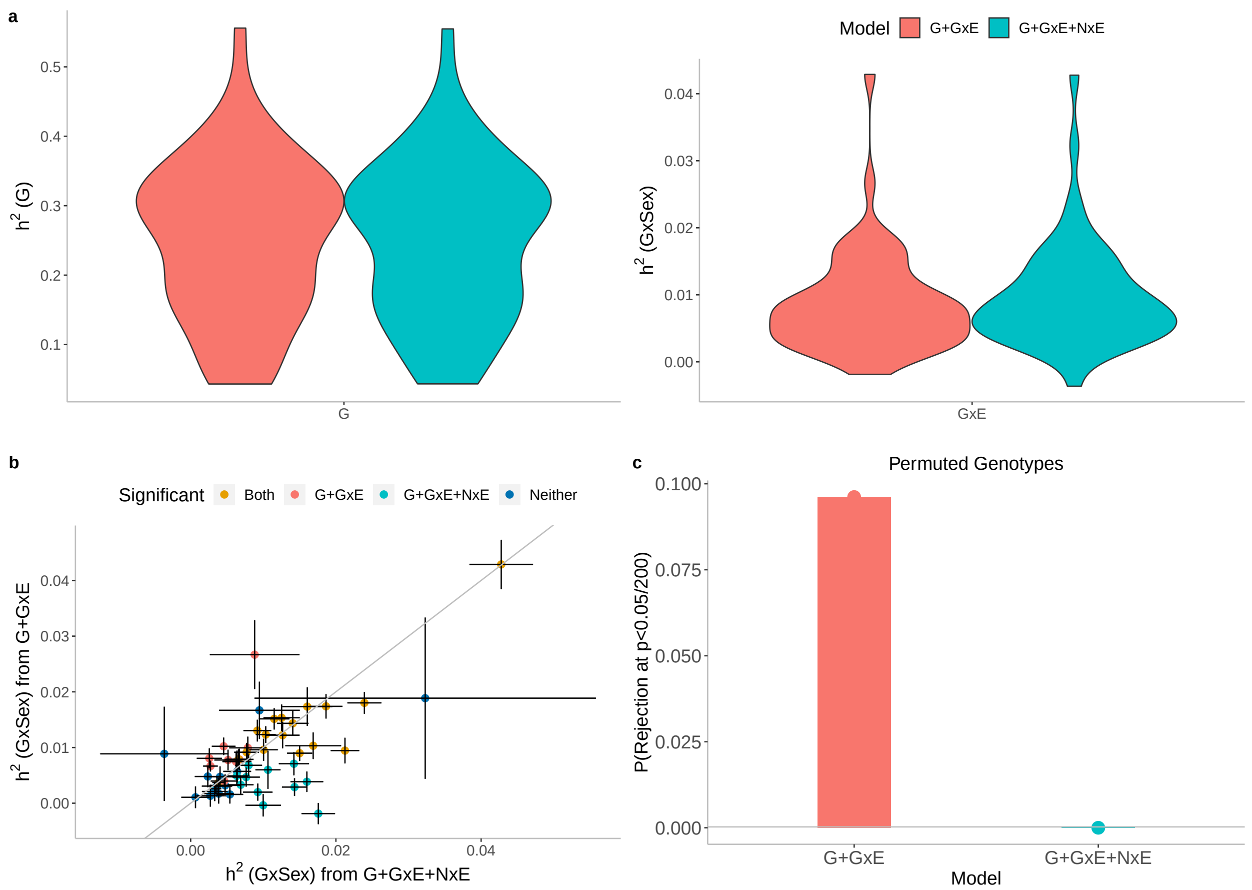


Figure S6: **Effect of Noise heterogeneity (NxE) on estimates of heritability associated with GxSex across 50 quantitative phenotypes in UKBB**. Model G+GxE refers to a model with additive and gene-by-environment interaction components. Model G+GxE+NxE refers to a model with additive, gene-by-environment interaction, and environmental heterogeneity (noise-by-environment interaction ) components. **a**) We run GENIE under G+GxE and G+GxE+NxE models to assess the effect of fitting an NxE component on the GxE and additive heritability estimates. **b**) Comparison of GxE heritability estimates obtained from GENIE under a G+GxE+NxE model (x-axis) to a G+GxE model (y-axis). Black error bars mark ± standard errors centered on the estimated GxE heritability. The color of the dots indicates whether estimates of GxE heritability are significant under each model. **c**) We performed permutation analyses by randomly shuffling the genotypes while preserving the trait-E relationship and applied GENIE in each setting under G+GxE and G+GxE+NxE models. We report the fraction of rejections (p-value of a test of the null hypothesis of zero GxE heritability $< \frac{0.05}{200}$ that accounts for the number of phenotypes tested) over 50 UKBB phenotypes.

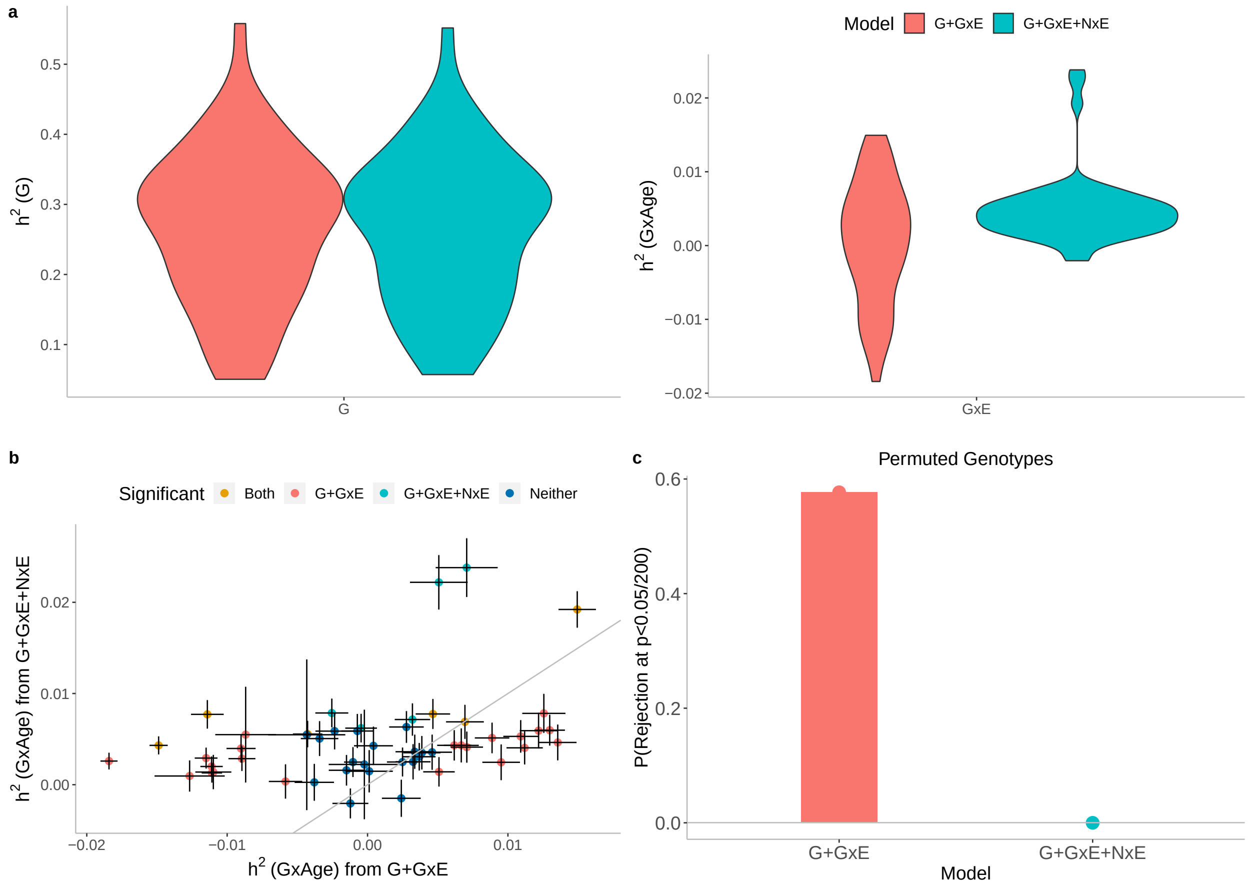


Figure S7: **Effect of Noise heterogeneity (NxE) on estimates of heritability associated with GxAge across 50 quantitative phenotypes in UKBB**. Model G+GxE refers to a model with additive and gene-by-environment interaction components. Model G+GxE+NxE refers to a model with additive, gene-by-environment interaction and environmental heterogeneity (noise-by-environment interaction ) components. **a**) We run GENIE under G+GxE and G+GxE+NxE models to assess the effect of fitting an NxE component on the GxE and additive heritability estimates. **b**) Comparison of GxE heritability estimates obtained from GENIE under a G+GxE+NxE model (x-axis) to a G+GxE model (y-axis). Black error bars mark ± standard errors centered on the estimated GxE heritability. Color of the dots indicate whether estimates of GxE heritability are significant under each model. **c**) We performed permutation analyses by randomly shuffling the genotypes while preserving the trait-E relationship and applied GENIE in each setting under G+GxE and G+GxE+NxE models. We report the fraction of rejections (p-value of a test of the null hypothesis of zero GxE heritability $< \frac{0.05}{200}$ that accounts for the number of phenotypes tested) over 50 UKBB phenotypes.

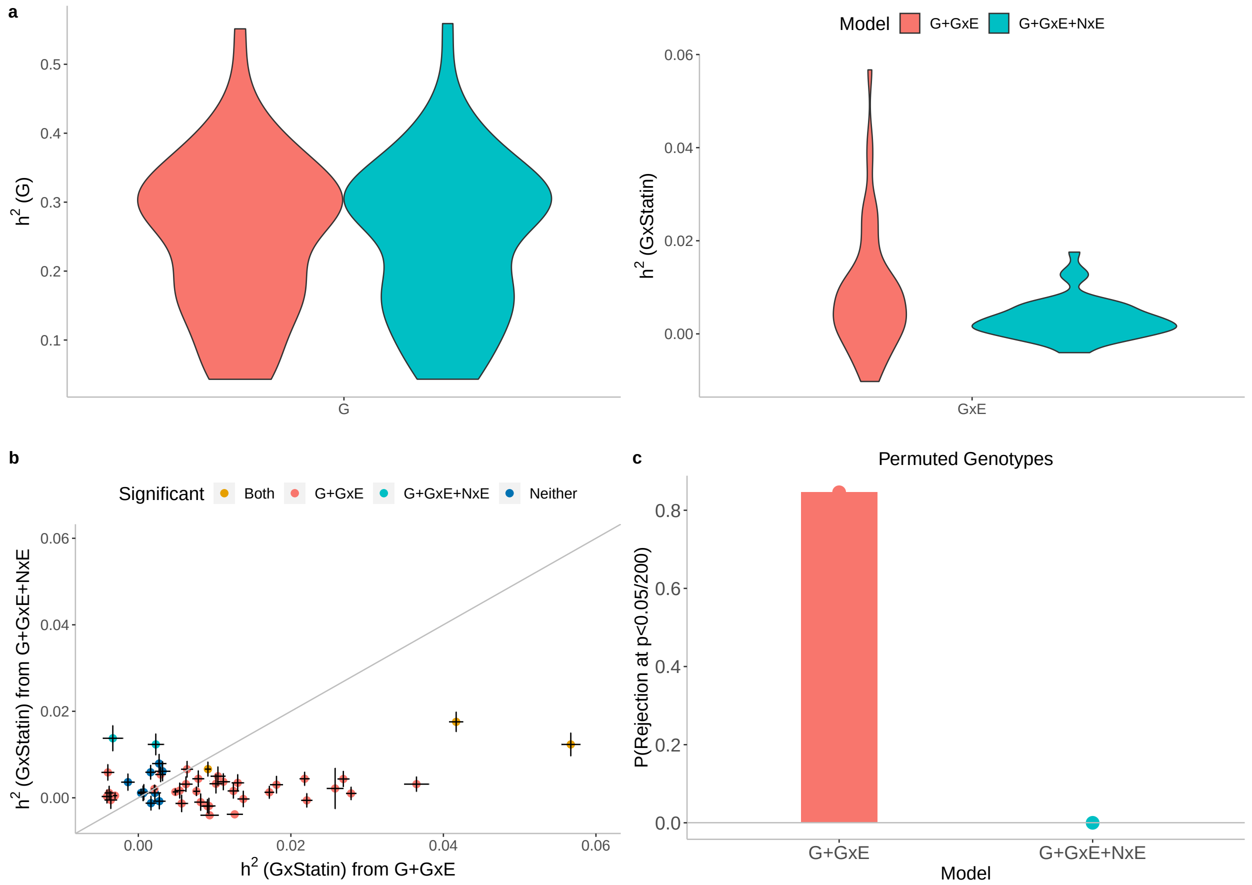


Figure S8: **Effect of Noise heterogeneity (NxE) on estimates of heritability associated with GxStatin across 50 quantitative phenotypes in UKBB**. Model G+GxE refers to a model with additive and gene-by-environment interaction components. Model G+GxE+NxE refers to a model with additive, gene-by-environment interaction and environmental heterogeneity (noise-by-environment interaction ) components. **a**) We run GENIE under G+GxE and G+GxE+NxE models to assess the effect of fitting an NxE component on the GxE and additive heritability estimates. **b**) Comparison of GxE heritability estimates obtained from GENIE under a G+GxE+NxE model (x-axis) to a G+GxE model (y-axis). Black error bars mark ± standard errors centered on the estimated GxE heritability. Color of the dots indicate whether estimates of GxE heritability are significant under each model. **c**) We performed permutation analyses by randomly shuffling the genotypes while preserving the trait-E relationship and applied GENIE in each setting under G+GxE and G+GxE+NxE models. We report the fraction of rejections (p-value of a test of the null hypothesis of zero GxE heritability $< \frac{0.05}{200}$ that accounts for the number of phenotypes tested) over 50 UKBB phenotypes.

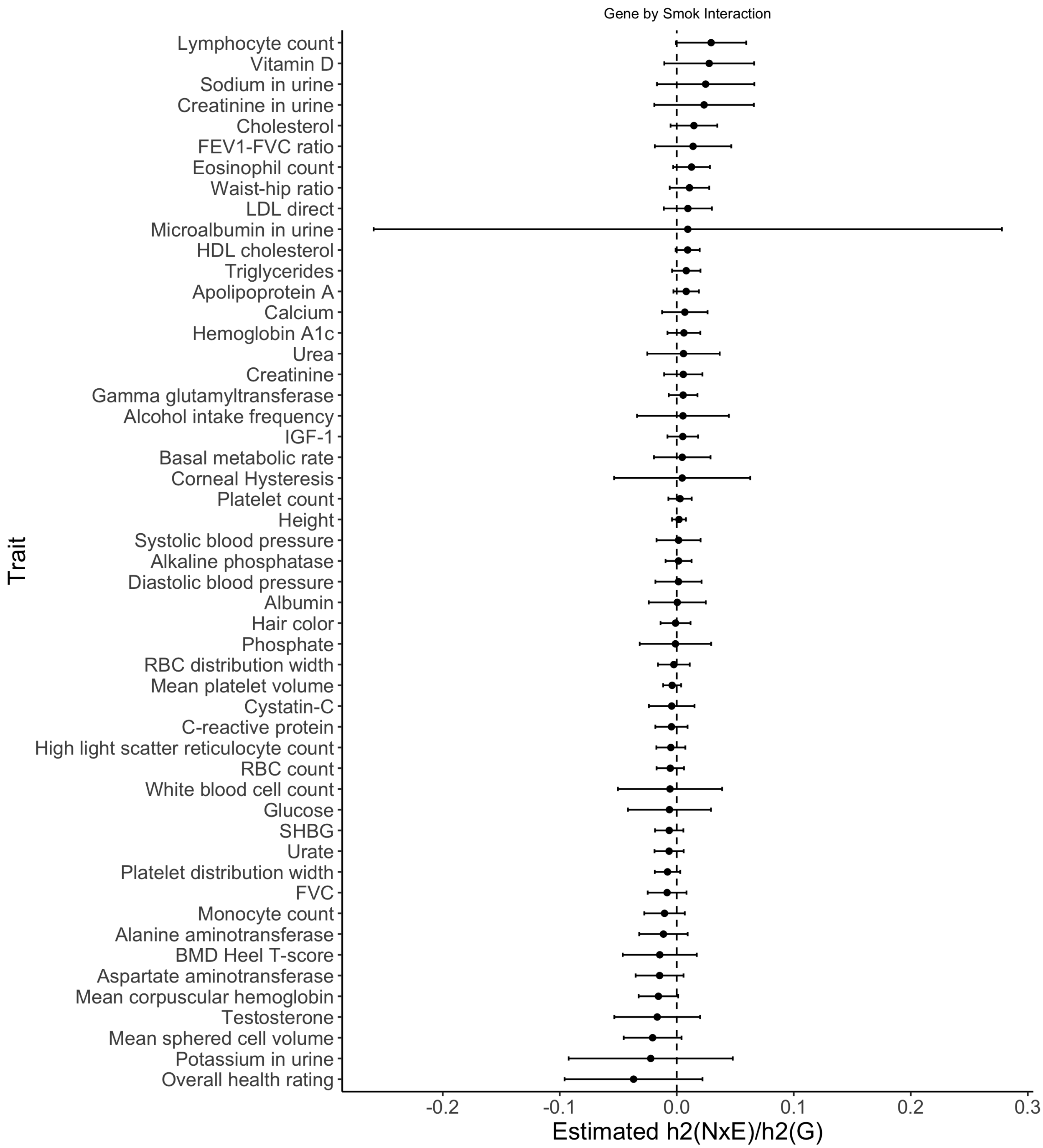


Figure S9: **Estimated ratio of variance attributed to noise heterogeneity over additive heritability for Smoking**. Black error bars mark ±2 standard errors centered on the estimated ratio.

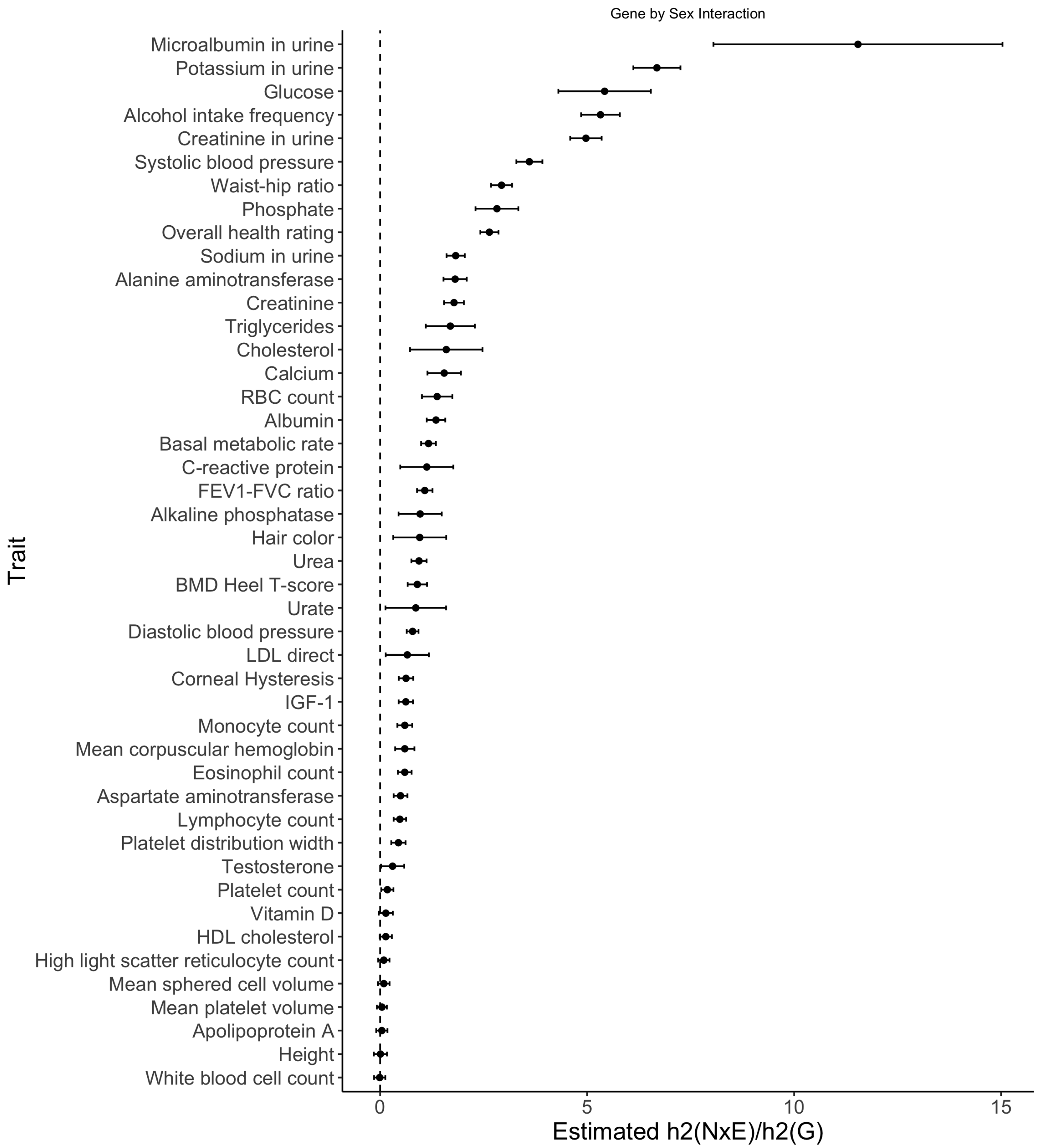


Figure S10: **Estimated ratio of variance attributed to noise heterogeneity over additive heritability for Sex**. Black error bars mark $\pm 2$ standard errors centered on the estimated ratio.

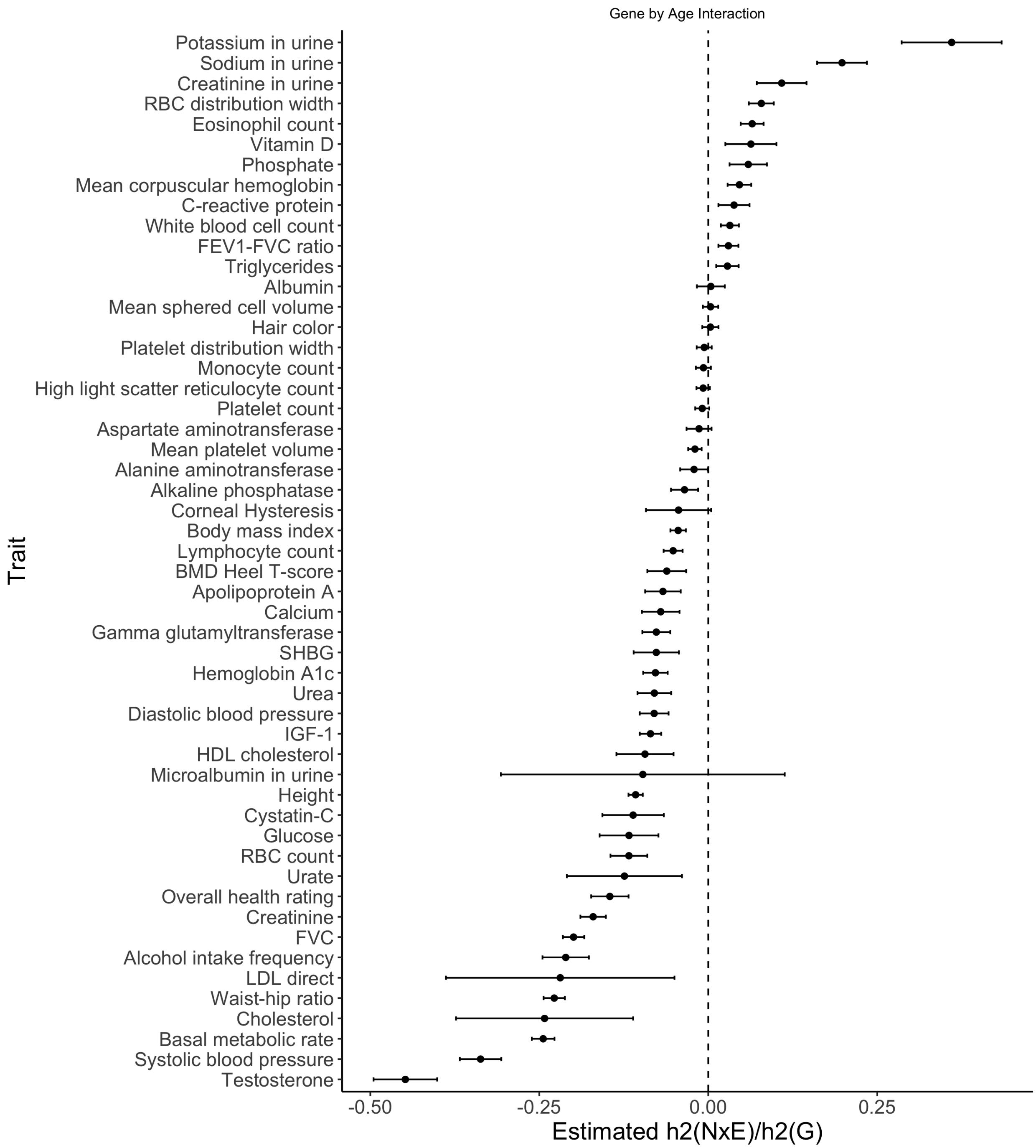


Figure S11: **Estimated ratio of variance attributed to noise heterogeneity over additive heritability for Age**. Black error bars mark ±2 standard errors centered on the estimated ratio.

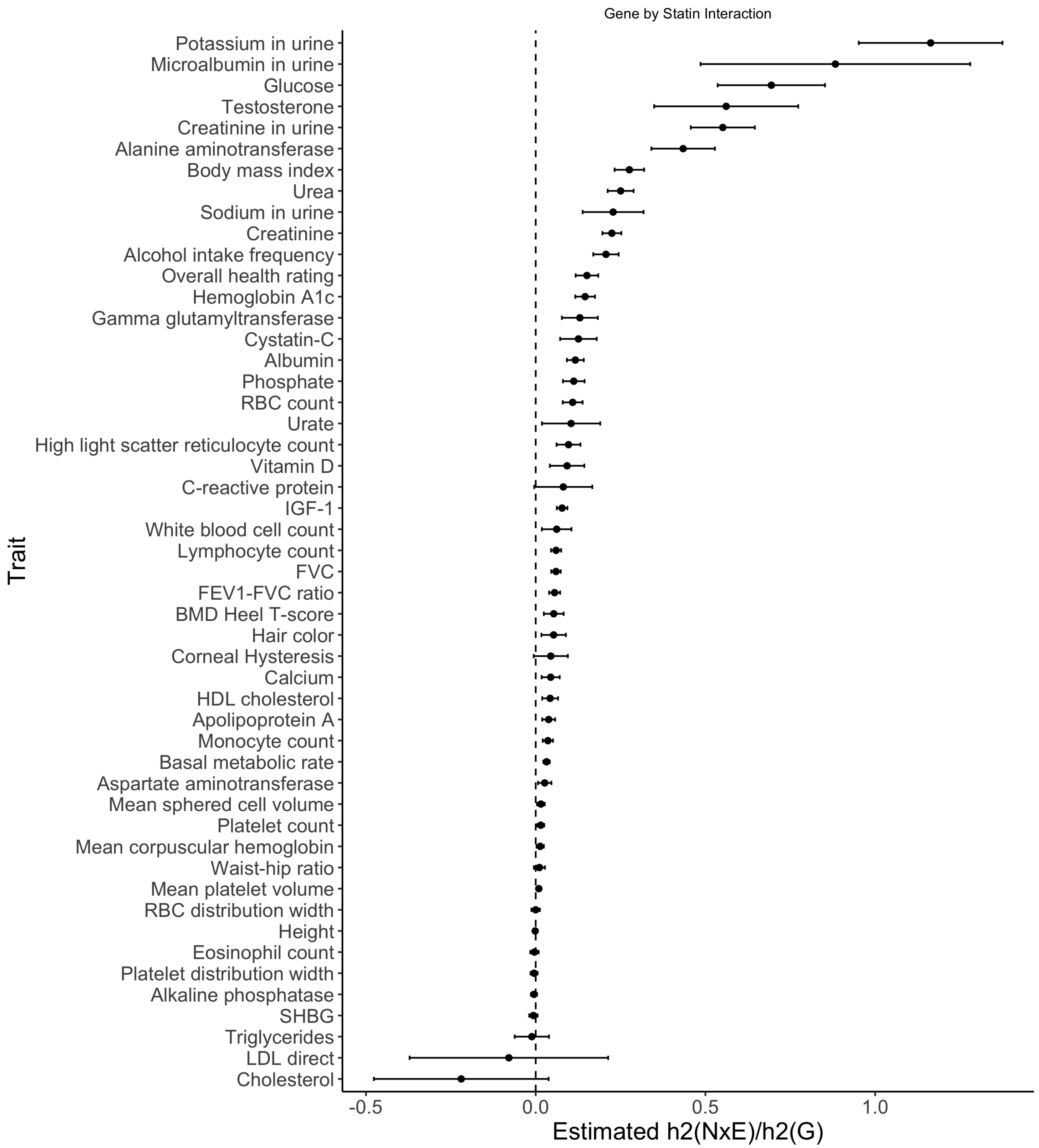


Figure S12: **Estimated ratio of variance attributed to noise heterogeneity over additive heritability for Statin usage**. Black error bars mark ±2 standard errors centered on the estimated ratio.

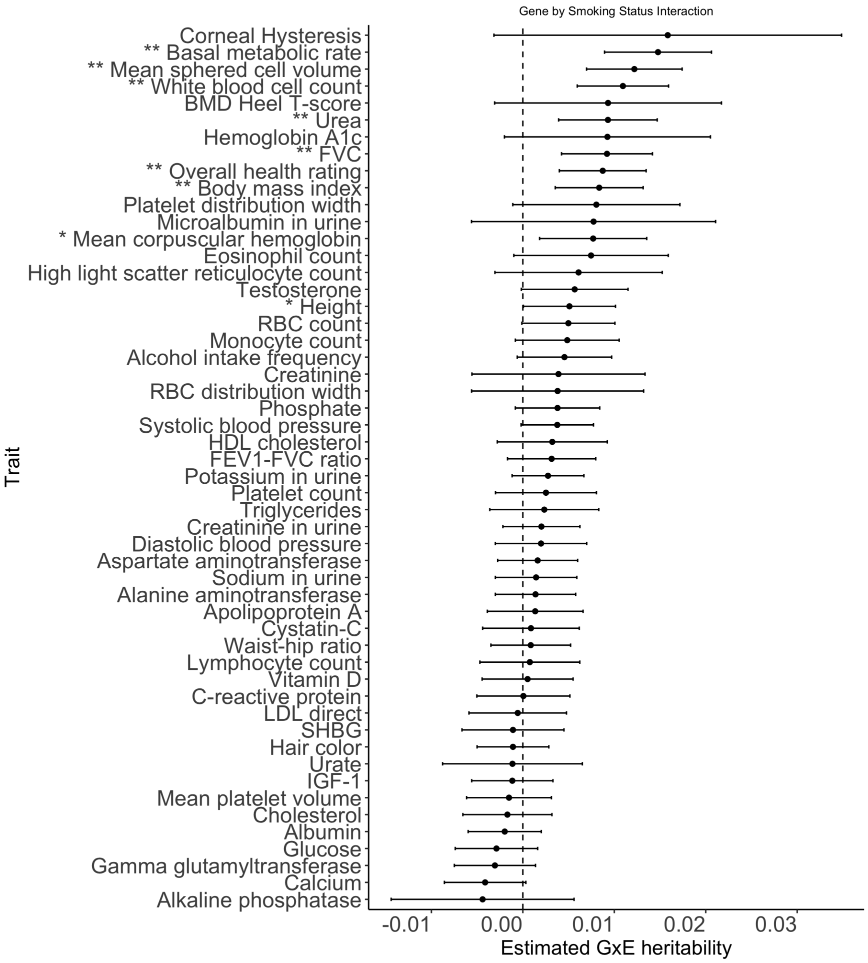


Figure S13: **GxSmoking across phenotypes in UK Biobank with the environmental variable coded as binary.** Our model includes the environmental variable as a fixed effect and accounts for environmental heterogeneity. Black error bars mark ±2 standard errors. The asterisk and double asterisk correspond to the nominal $p < 0.05$ and $p < 0.05/200$ respectively.

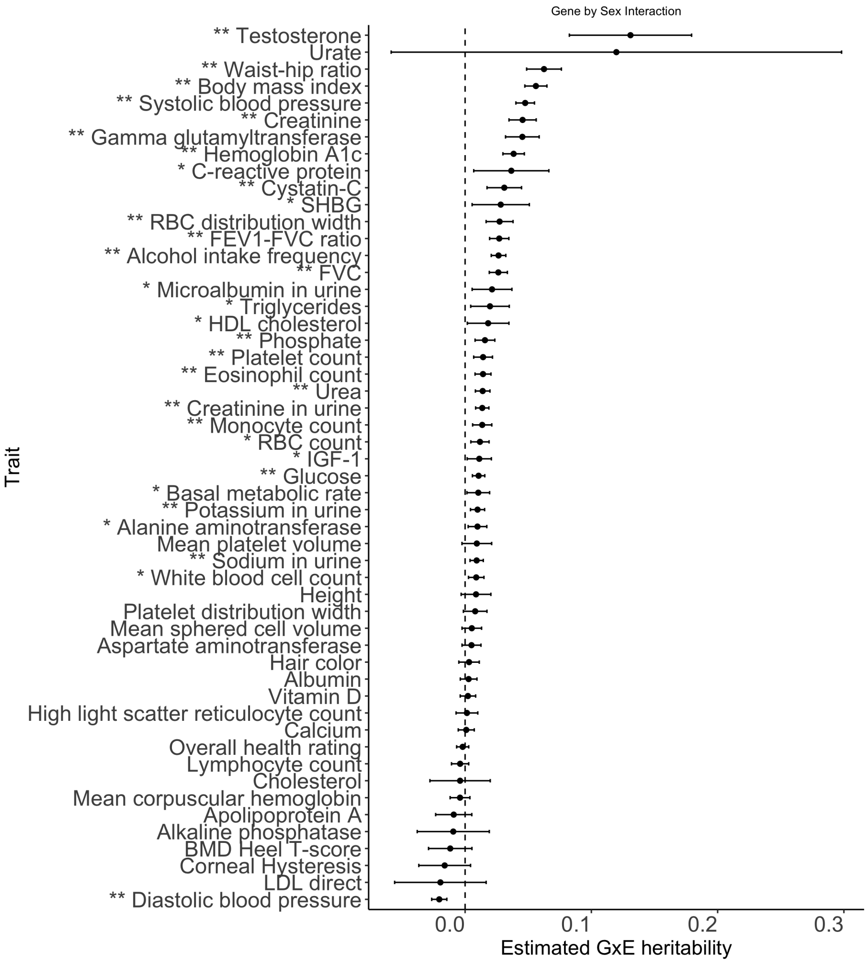


Figure S14: **GxSex across phenotypes in UK Biobank with the environmental variable coded as binary.** Our model includes the environmental variable as a fixed effect and accounts for environmental heterogeneity. Black error bars mark $\pm 2$ standard errors. The asterisk and double asterisk correspond to the nominal $p < 0.05$ and $p < 0.05/50$ respectively.

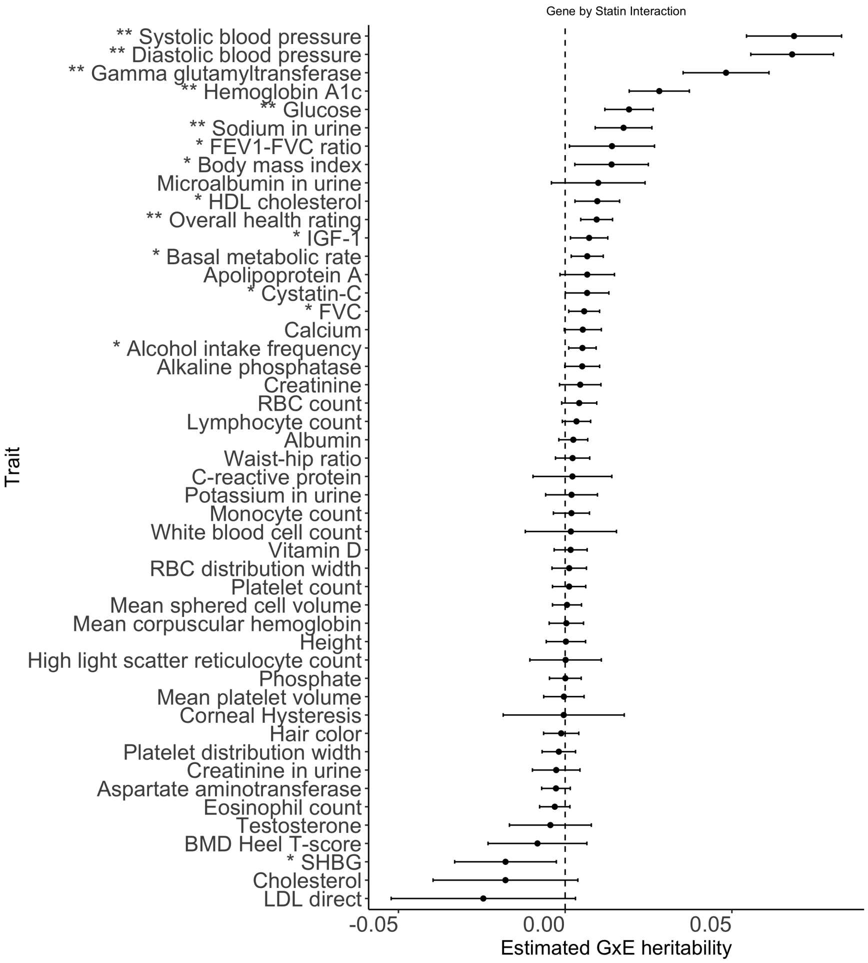


Figure S15: **GxStatin across phenotypes in UK Biobank with the environmental variable coded as binary.** Our model includes the environmental variable as a fixed effect and accounts for environmental heterogeneity. Black error bars mark $\pm 2$ standard errors. The asterisk and double asterisk correspond to the nominal $p < 0.05$ and $p < 0.05/50$, respectively.

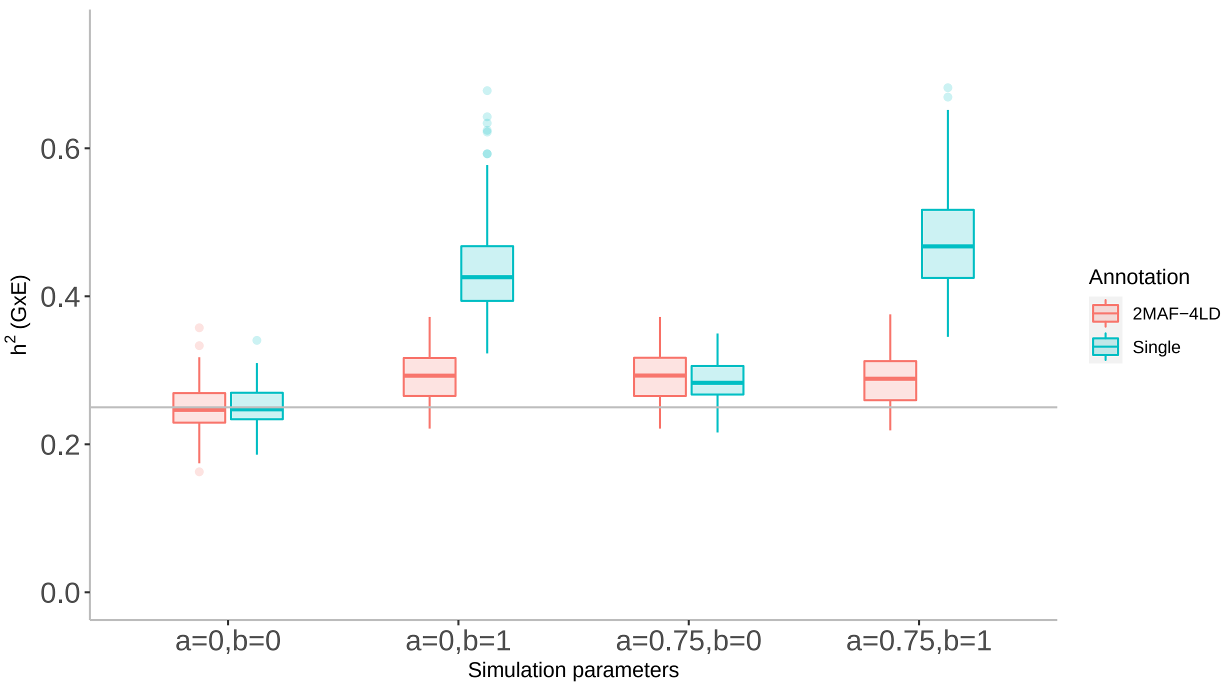


Figure S16: **Effect of MAF-LD partitioning on estimated GxE heritability in simulation.** We assessed the effect of MAF-LD partitioning on estimates of $h^2_{gxe}$ in simulations. We ran GENIE in two settings: 1) fitting a model with a single additive and a single GxE variance component, 2) fitting a model with eight additive and eight GxE components defined based on four LD annotations (quartiles of LD scores) and two MAF annotations. we simulated phenotypes with GxE effects and G effects from a subset of $N = 40k$ individuals genotyped at array SNPs $M = 459,792$ by varying the coupling of MAF with effect size $(a)$ and the effect of local LD on effect size (b) (see Supplementary note S2 for details). Here we have $h^2_g = h^2_{gxe} = 0.25$,$h^2_{nxe} = 0.05$, and all SNPs are causal for both additive and GxE effects. Each box plot represents estimates from 100 simulations.

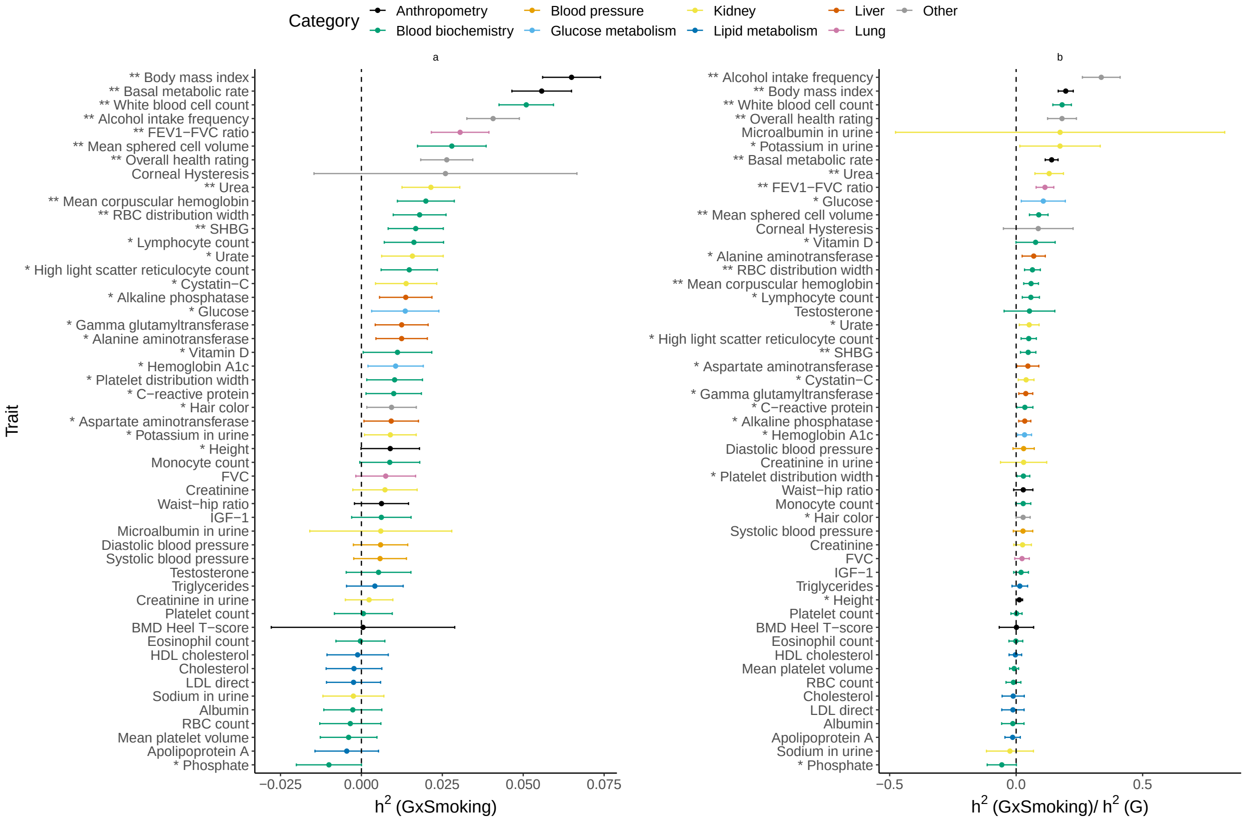


Figure S17: **GxSmoking across phenotypes from imputed SNPs in UK Biobank by MAF-LD partitioning.** Our model includes the environmental variable as a fixed effect and accounts for environmental heterogeneity. Black error bars mark $\pm 2$ standard errors. The asterisk and double asterisk correspond to the nominal $p < 0.05$ and $p < 0.05/200$, respectively.

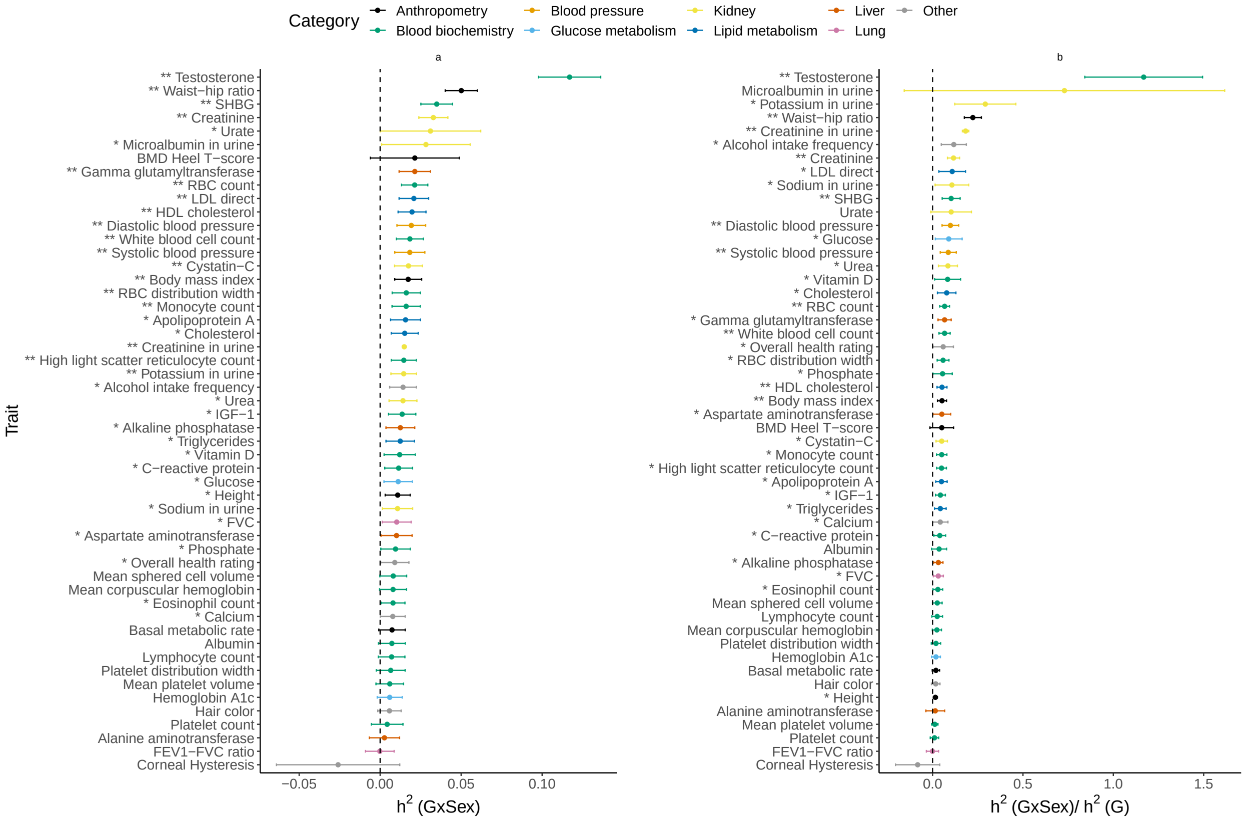


Figure S18: **GxSex across phenotypes from imputed SNPs in UK Biobank by MAF-LD partitioning.** Our model includes the environmental variable as a fixed effect and accounts for environmental heterogeneity. Error bars mark ±2 standard errors. The asterisk and double asterisk correspond to the nominal $p < 0.05$ and $p < 0.05/200$, respectively.

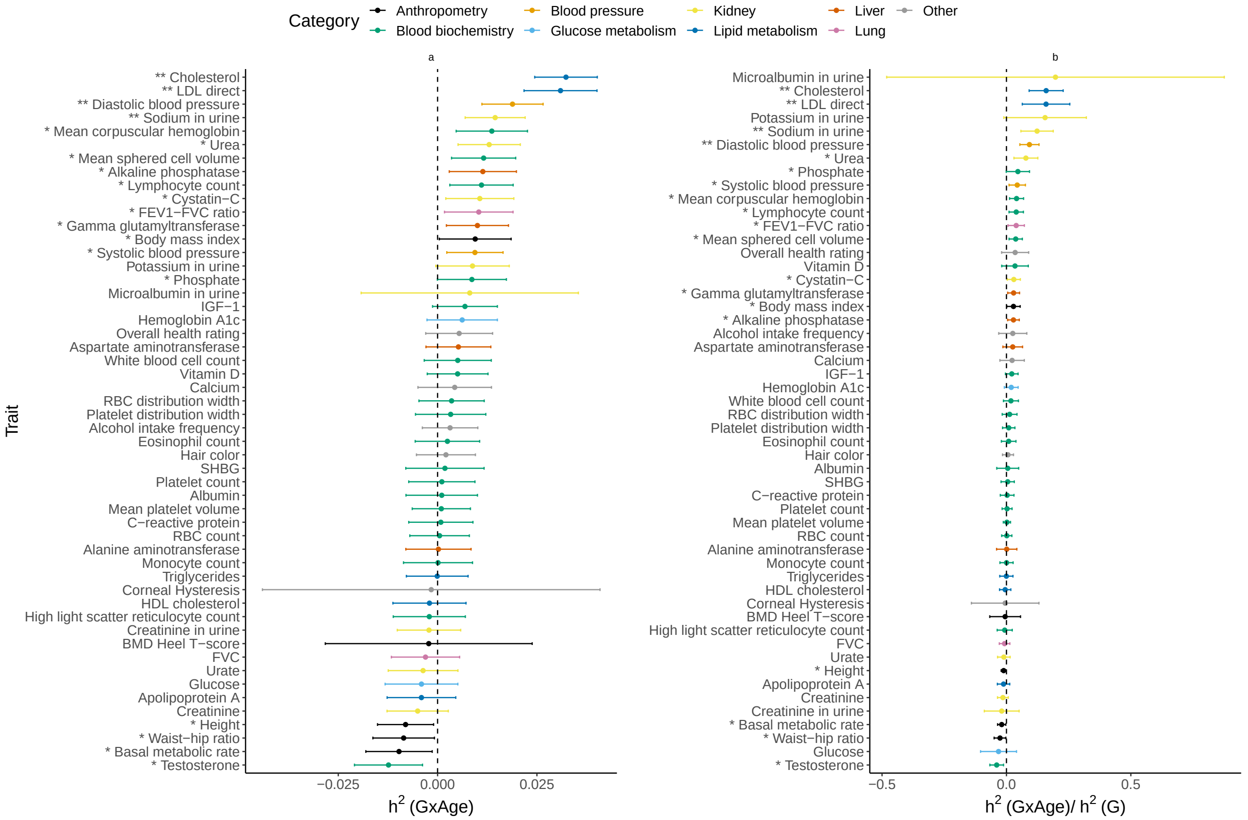


Figure S19: **GxAge across phenotypes from imputed SNPs in UK Biobank by MAF-LD partitioning.** Our model includes the environmental variable as a fixed effect and accounts for environmental heterogeneity. Error bars mark ±2 standard errors. The asterisk and double asterisk correspond to the nominal $p < 0.05$ and $p < 0.05/200$, respectively.

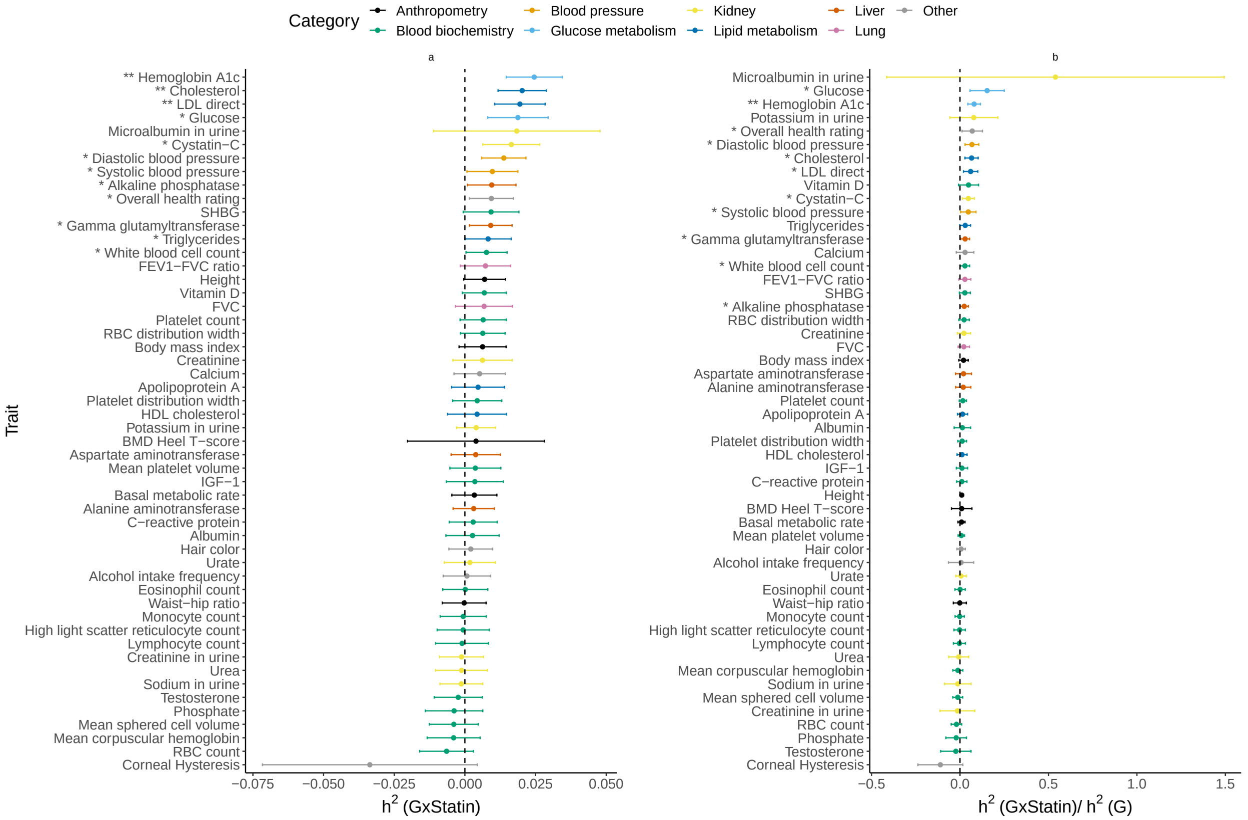


Figure S20: **GxStatin across phenotypes from imputed SNPs in UK Biobank by MAF-LD partitioning.** Our model includes the environmental variable as a fixed effect and accounts for environmental heterogeneity. Error bars mark ±2 standard errors. The asterisk and double asterisk correspond to the nominal $p < 0.05$ and $p < 0.05/200$, respectively.

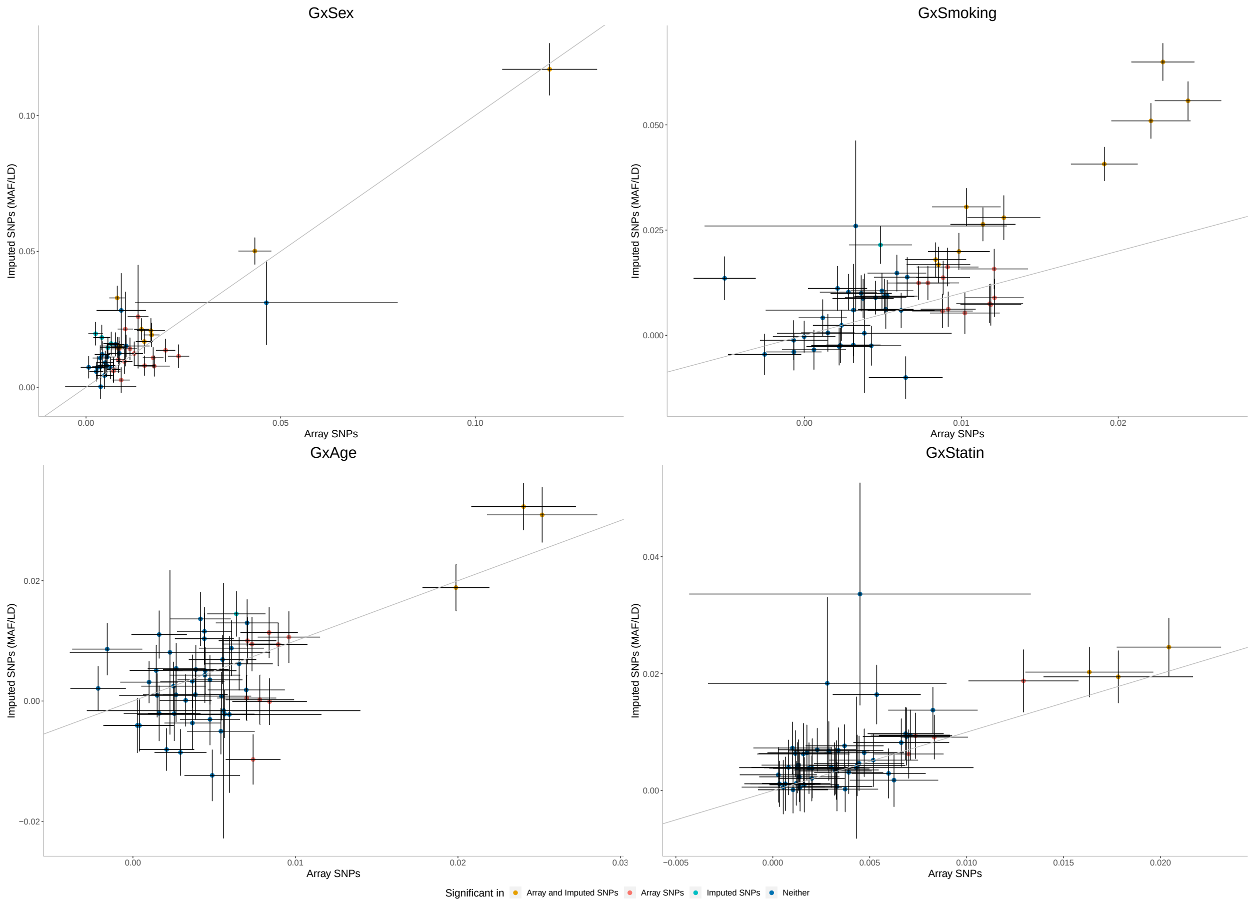


Figure S21: **Comparing GxSex, GxSmoking, GxAge, and GxStatin estimates from imputed SNPs (MAF$\geq$ 0.1%) and array SNPs (MAF$\geq$ 1%).** In this analysis, we applied GENIE to imputed SNPs with MAF/LD stratification and array SNPs with a single component. Black error bars mark $\pm 2$ standard errors. The asterisk and double asterisk correspond to the nominal $p < 0.05$ and $p < 0.05/200$ respectively. Color of the dots indicate whether estimates of GxE heritability are significant under each model.

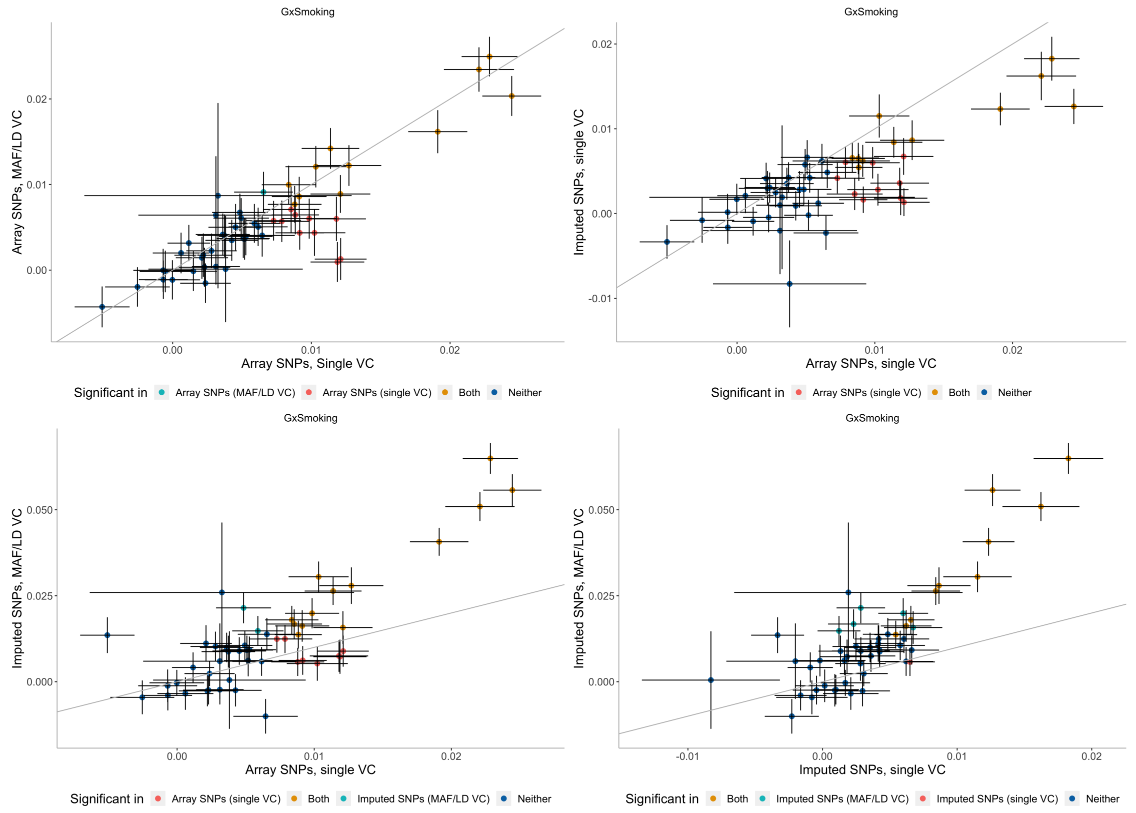


Figure S22: **Effect of MAF-LD partitioning on estimated GxE heritability.** We assessed the effect of MAF-LD partitioning on estimates of $h^2_{gxSmoking}$ from array SNPs and imputed SNPs. We ran GENIE in two settings: 1) fitting a model with a single additive and a single GxE variance component, 2) fitting a model with eight additive and eight GxE components defined based on four LD annotations (quartiles of LD scores) and two MAF annotations. Black error bars mark $\pm 2$ standard errors centered on the estimates of $h^2_{gxSmoking}$. Color of the dots indicate whether estimates of $h^2_{gxSmoking}$ are significant under each model.

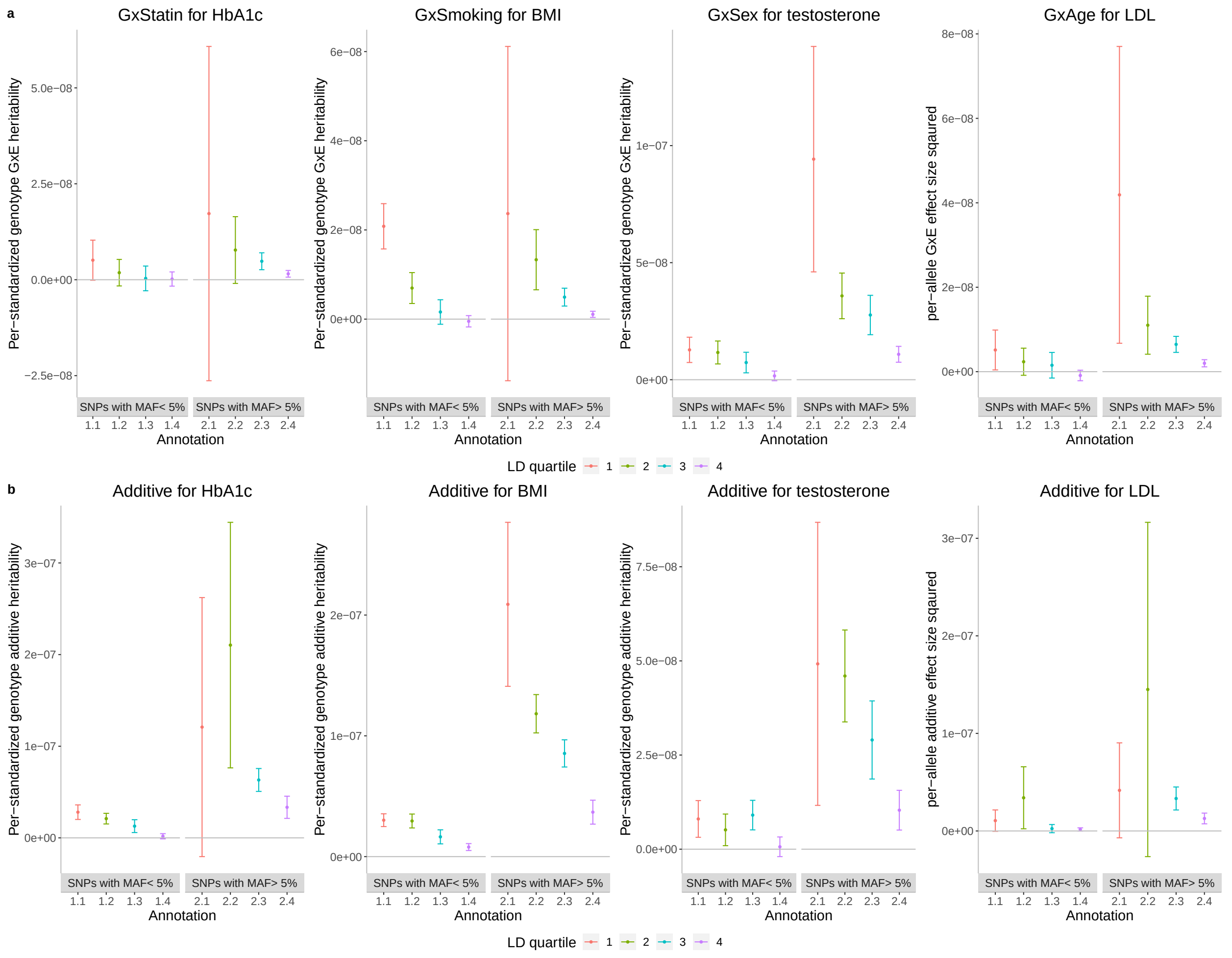


Figure S23: **Per-standardized genotype GxE and additive heritability as a function of MAF and LD.** **a**) The per-standardized genotype GxE heritability for four selected pairs of traits and environments (trait-E pairs). **b**) The per-allele additive heritability for the same trait-E pairs. The x-axis corresponds to MAF-LD annotations where annotation $i.j$ includes SNPs in MAF bin $i$ and LD quartile $j$ where MAF bin 1 and MAF bin 2 correspond to SNPs with MAF $\leq 5\%$ and MAF $> 5\%$ respectively while the first quartile of LD-scores correspond to SNPs with the lowest LD-scores respectively). The y-axis shows the per-standardized genotype GxE (or additive) heritability defined as $\frac{h_k^2}{2M_k}$ where $h_k^2$ is the GxE (or additive) heritability attributed to bin $k$, $M_k$ is the number of SNPs in bin $k$. Error bars mark $\pm 2$ standard errors centered on the estimated effect sizes.

[40] Doug Speed and David Balding. Better estimation of snp heritability from summary statistics provides a new understanding of the genetic architecture of complex traits. *bioRxiv*, page 284976, 2018.

[41] Steven Gazal, Hilary K Finucane, Nicholas A Furlotte, Po-Ru Loh, Pier Francesco Palamara, Xuanyao Liu, Armin Schoech, Brendan Bulik-Sullivan, Benjamin M Neale, Alexander Gusev, et al. Linkage disequilibrium–dependent architecture of human complex traits shows action of negative selection. *Nature genetics*, 49(10):1421, 2017.

[42] Ali Pazokitoroudi, Yue Wu, Kathryn S. Burch, Kangcheng Hou, Aaron Zhou, B. Pasaniuc, and S. Sankararaman. Efficient variance components analysis across millions of genomes. *Nature Communications*, 11, 2020.
